# YAP1 is a key regulator of EWS::FLI1-dependent malignant transformation upon IGF-1 mediated reprogramming of bone mesenchymal stem cells

**DOI:** 10.1101/2024.07.15.603565

**Authors:** Rahil Noorizadeh, Barbara Sax, Tahereh Javaheri, Branka Radic Sarikas, Valerie Fock, Maximilian Kauer, Aleksandr Bykov, Veveeyan Suresh, Michaela Schlederer, Lukas Kenner, Gerhard Weber, Wolfgang Mikulits, Florian Halbritter, Richard Moriggl, Heinrich Kovar

## Abstract

Ewing sarcoma (EwS) is an aggressive cancer of adolescents in need of effective treatments. Insulin like growth factor (IGF) 1 was previously reported an autocrine growth factor for EwS, but only 10% of patients responded to IGF-1 receptor blockade. Although presumed to originate from mesenchymal progenitors during bone development, targeting of the EwS driver oncogene *EWS::FLI1* to the mesenchymal lineage in a conditional mouse model did not result in tumor formation but led to skeletal malformations and perinatal death. We report that transient exposure to IGF-1 concentrations mimicking serum levels during puberty reprogrammed limb-derived mesenchymal cells of *EWS::FLI1*-mutant mice to stable transformation and tumorigenicity. We identified a modular mechanism of IGF-1-driven tumor promotion in the early steps of EwS pathogenesis, in which Yap1 plays a central role. Pharmacologic Yap1/Tead inhibition reversed the transformed phenotype of EWS::FLI1 expressing cells. Our data provide a rationale for combined IGF-1R and YAP/TEAD inhibition in the treatment of EwS patients.

**Graphical Abstract:** 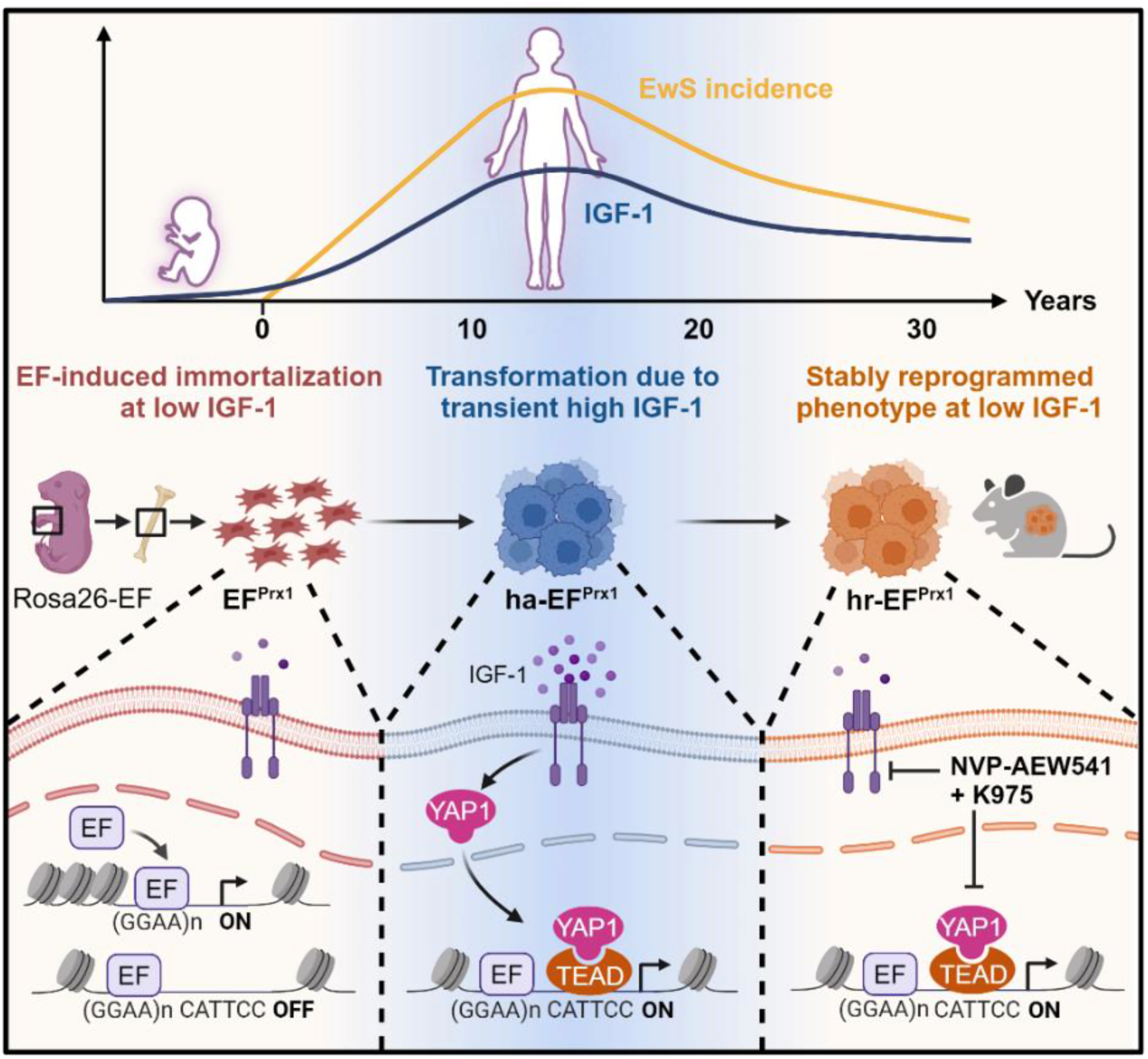

## Introduction

Ewing sarcoma (EwS) is a highly aggressive bone and soft-tissue cancer with a peak incidence during puberty ^1^. The 5-year disease-free survival rate of patients with localized EwS is about 70-80%, while only 20-30% of patients with metastases survive beyond 5 years despite intensive multi-modal treatment. Therefore, there is an urgent unmet need for more efficient therapeutic approaches in this group of patients. However, progress in the identification and validation of novel treatment options is limited by a paucity in preclinical *in vivo* models faithfully recapitulating the human disease ^2,3^.

EwS is characterized by a chromosomal translocation rearranging the *EWSR1* gene on chromosome 22 with an ETS transcription factor family gene, most commonly *FLI1* ^4^. The resulting gene fusion encodes the oncogenic transcription factor EWS::FLI1 (EF), targeting the aberrant expression and processing of a large number of genes involved in cellular proliferation and transformation ^1^. Previous studies have shown that EF acts as a unique pioneer factor at GGAA repeat sites, mediating a transition from closed to open chromatin and establishing an active enhancer state leading to the aberrant activation of many EwS hallmark genes ^1^. Despite extensive knowledge on EF-driven gene regulatory mechanisms, the generation of a genetic mouse model for EwS remains challenging due to toxicity of EF to most cell types and body tissues ^2^. Early studies in mouse fibroblasts revealed dependence of EF-driven transformation on the expression of the insulin-like growth factor-1 receptor (IGF-1R) ^5^. In addition, autocrine production of IGF-1 was demonstrated to depend on EF expression in EwS ^6,7^. In fact, evidence suggests that EF is creating a cellular environment conducive for IGF-1R signaling. EF directly activates the *IGF-1* promoter ^6^ and simultaneously suppresses IGF-1R signaling-modulatory pathways through downregulation of several IGF-1 and IGF-1R targeting microRNAs ^8^. In addition, EF directly suppresses transcription as well as pappalysine-1 mediated degradation of negative regulatory IGF-1-binding proteins (IGFBPs) ^9,10^. Consequently, experimental silencing of EF expression resulted in impaired IGF-1R signaling ^11^. Vice versa, blocking IGF-1 signaling by IGF-1R antagonists, either antibodies or small molecule inhibitors, inhibited EwS growth in athymic mice ^12^. However, only 10-14% of patients responded to IGF-1R-directed therapy in clinical trials ^13–15^, at least in part due to upregulation of the closely related and interconnected Insulin receptor signaling pathway, where both receptors form heterodimers ^16^. Collectively, these results demonstrate dependence of sustained EwS growth on EF-driven autocrine IGF-1/Insulin signaling, but the status and role of this pathway at early stages of EwS pathogenesis remain unknown.

Identification of the EwS cell of origin and its developmental stage is crucial to understand EwS initiation and progression. Neural crest-and mesoderm-derived mesenchymal stem cell (MSC) compartments are widely considered candidate progenitor cell types for EwS ^1^. Consistent with this hypothesis, targeting transgenic EF expression to the mesenchymal lineage in p53 knockout mice accelerated sarcoma formation ^17^, but both germline and somatic p53 alterations are rare in human EwS with an incidence below 10% ^18–20^.

We took a similar approach to restrict EF expression to the mesenchymal lineage during endochondral bone formation and confirm that early expression of EF during embryogenesis is associated with severe skeletal malformations due to differentiation arrest at an early chondrocytic stage leading to death of progenies a few hours after birth. As EwS incidence in humans peaks during the second decade of life, a developmental period in which several growth-promoting hormones are highly enriched in the bone microenvironment ^21,22^, we hypothesized that this endocrine milieu may play a tumor promoting role in EwS pathogenesis. Interaction between tumor cells, tumor-derived humoral factors, and the bone marrow in the bone niche has been shown to be essential for bone tumor initiation and progression ^23,24^. We focused on IGF-1, which is known for its anti-apoptotic activity through upregulation of Bcl2 family genes ^25^. In the bone niche, IGF-1 is expressed from osteocytes and osteoblasts in response to mechanical load stimulating chondrocyte differentiation and bone growth ^26,27^. Here, we provide evidence that mouse embryonal EF-expressing limb-derived mesenchymal stem cell-like cells (MSCLC) can be fully and stably transformed if transiently exposed to human pubertal serum IGF-1 levels, a condition that EwS precursor cells may experience in the bone niche during adolescence. We describe a modular reprogramming mechanism resulting in the activation of EwS hallmark genes with previously documented roles in EwS growth and survival, and identify a key functional role for the transcriptional co-regulator Yap1 and its Tead transcription factor effectors. Our study provides a mechanism underlying the trajectories from EF-driven immortalization to full transformation caused by the activation of bone developmental signaling cues.

## Results

### Embryonal expression of EF in the mesenchymal lineage impairs endochondral bone formation

We restricted EF expression to the mesenchymal lineage of endochondral bone formation by crossing a mouse line carrying a *loxP-STOP-loxP*-HA-tagged EF cassette knocked in the *Rosa26* ^28^ to a Prx1-Cre transgenic line ^29^. Mesenchymal targeting of Cre recombinase led to deletion of the *STOP* cassette allowing for *Rosa26* promoter-driven expression of HA-tagged EF in the developmental limb mesenchyme starting from embryonal day E9.5 ^29^ (Figure 1A, Figure S1). The resulting mice (hereafter referred to as EF^Prx1^) displayed severe skeletal malformations, including polydactyly, short limbs, craniofacial malformations, and lack of rib cage closure. They died within ∼12 hours after birth. The aberrant skeletal phenotype was already evident at an early stage of embryonic development on embryonic day 15.5 (E15.5) (Figure 1B). We performed a skeletal analysis with Alcian Blue/Alizarin Red staining at E18.5, which revealed an absence of calcification in EF^Prx1^ limbs, sternum, and head bone calvaria (Figure 1C). Histological analysis of E18.5 EF^Prx1^ limbs identified only condensed cartilaginous elements with a lack of a hypertrophic zone (Figure 1D). Consistent with this finding and different from parental Prx1-Cre mice used as wild-type controls, RNA in-situ hybridization analysis was negative for chondrocyte markers such as Indian hedgehog signaling molecule (Ihh) and collagen type 10 alpha 1 chain (Col10a1).

**Fig. 1.**
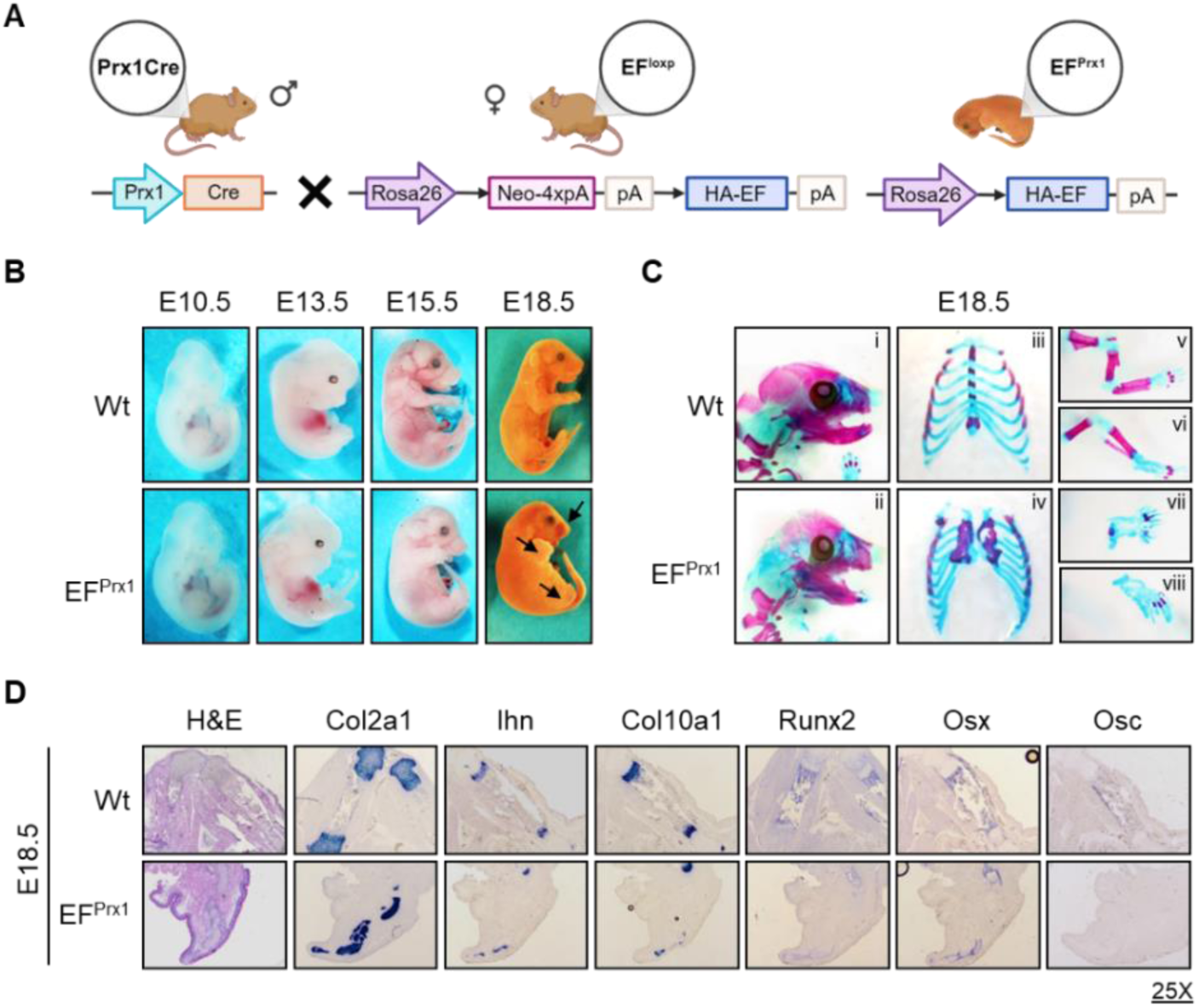
Early EF expression in MSCLC during embryogenesis causes endochondral bone formation arrest. **A** Schematic drawing of EF^Prx1^ mice generation by crossing Prx1-Cre mice with Rosa26 loxP-STOP-loxP-HA-EF (EF^loxp^) mice. Black arrows: loxP sites. Hemagglutinin tag: HA, EWS::FLI1: EF. **B** EF^Prx1^ mouse embryos exhibit multiple abnormalities of the limbs and the cranium (blue dashed arrows). **C** Skeletal preparations and analysis with Alcian Blue/Alizarin Red staining of E18.5 Wt and EF^Prx1^ embryos. Blue staining marks cartilage elements, whereas pink color indicates calcified bone. EF^Prx1^ embryos had defects in the development of craniofacial (i,ii), the sternum (iii, iv), forelimbs (v, vii), and hindlimbs (vi,viii). **D** Hematoxylin and Eosin staining (H&E) of limb sections of E18.5 Wt and EF^Prx1^ mouse embryos (a), and non-radioactive RNA in-situ hybridization for expression of markers specific to different chondrocytic (Col2a1, Col10a1, Ihh) and osteoblastic (Runx2, Osx, Col1a1, Osc) maturation stages (25x magnification).

Additionally, EF^Prx1^ limbs were also deficient for osteoblast markers, including runt-related transcription factor 2 (Runx2), osterix (Osx), and osteocalcin (Osc) (Figure 1D). Together, these results indicate an EF-induced developmental differentiation arrest of MSCLC at an early chondrocytic stage during endochondral bone formation explaining the observed severe shortening of limbs and the lack of rib cage closure in EF^Prx1^mice (Figure 1C). The latter malformation contributed to the suffocation of newborn mice as a main reason for perinatal death paired with an inability to suckle milk due to craniofacial/calvaria malformation.

### Characterization of limb-derived mesenchymal stem cell-like cells from EF mutant mice

To more closely characterize the molecular underpinnings of the observed differentiation arrest, we isolated and propagated MSCLC from the limbs of two newborn EF^Prx1^ (#1 and #2) and wild-type Prx1-Cre mice (Wt). EF^Prx1^ MSCLCs stably expressed EF protein (Figure S2A) and were morphologically (Figure S2B) and immunophenotypically similar to Wt MSCLC, staining positive for mesenchymal markers CD90.2 and CD44 and negative for hematopoietic lineage markers CD45 and CD19 (Figure S2C). However, when incubated with the appropriate cytokine cocktails, EF^Prx1^ MSCLC retained chondrocytic differentiation potential but failed to differentiate into adipocytic and osteoblastic lineages, while Wt controls were able to differentiate into all three lineages (Figure S2D). RNA-seq analysis identified several anti-apoptotic *Bcl2a* family members among 2,669 genes significantly downregulated in EF^Prx1^ MSCLC as compared to Wt MSCLC (DESeq2; P_adj_ <= 0.05, |log_2_FC| >= log_2_(2)) ^30^ (Figure S3A and S3B; Tables S1 and S2). Among 1,610 genes that were found to be upregulated in EF^Prx1^ MSCLC in comparison to Wt MSCLC, a significant enrichment of previously published human EF signature gene sets was observed (hypergeometric tests with hypeR; KINSEY_TARGETS_OF_EWSR1_FLII_FUSION_UP: P_adj_ = 0.00032, RIGGI_EWING_SARCOMA_PROGENITOR_UP: P_adj_ = 0.068; Figure S3C and S3D; Table S3).

In EwS, EF is known to gain the unique neomorphic ability to bind and open microsatellite sequences consisting of multiple “GGAA” repeats, turning them into transcriptional enhancers ^1^. To monitor the effects of EF expression on global chromatin accessibility in our model, we applied the assay for transposase-accessible chromatin using sequencing (ATAC-seq)^31^ analysis in EF^Prx1^ #1 and #2 versus Wt MSCLC (Tables S1 and S4). 529 regions were found to have opened and 972 regions to have closed in EF^Prx1^ MSCLC compared to the Wt MSCLC (Figure 2A, Table S5). The majority (84.3%) of ATAC-seq peaks gained in EF^Prx1^ MSCLC concentrated at a distance of more than 10kb from the transcription start site (TSS) of the nearest gene, while 31.6% of peaks unaltered or lost in EF^Prx1^ MSCLC in comparison to Wt MSCLC localized in proximal regions (<10kb from TSS) (Figure 2B). In addition, we found an enrichment of GGAA microsatellites in differentially opened regions of EF^Prx1^ MSCLC (fGSEA; FDR = 0.005)^32^ (Figures 2C and 2D). Together, these data suggest that in mouse EF^Prx1^ MSCLC, EF retains its neomorphic activity epigenetically reprogramming GGAA microsatellite enhancers.

**Fig. 2.**
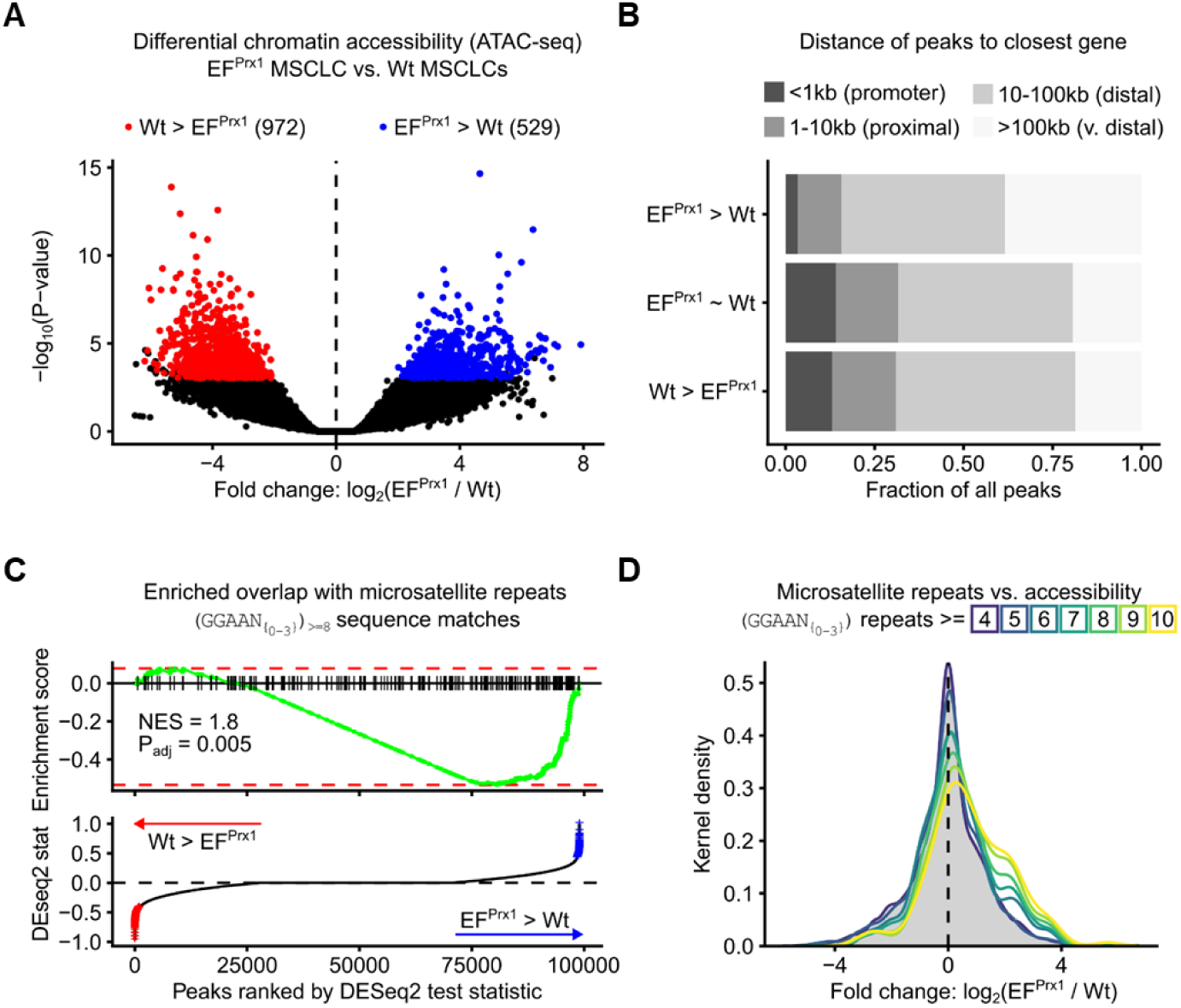
Chromatin accessibility is altered in EF^Prx1^ MSCLC. **A** Volcano plot of differentially accessible chromatin regions between EF^Prx1^ MSCLC and Wt MSCLC identified by DEseq2. The X-axis indicates the logarithms of the fold changes of individual genes. The Y-axis indicates the negative logarithm of their P-value to base 10 (DESeq2 ^30^; P_adj_<0.05, |log_2_FC|>log_2_(1.5); n(EF^Prx1^) = 4, n(Wt) = 3). See Tables S4, S5. **B** Distribution of differentially accessible peaks (from **A**) with respect to the distance from the transcription start site of the nearest gene compared to all regions without significant change (EF^Prx1^ MSCLC ≈ Wt MSCLC). **C** Fast Gene set enrichment analysis (fGSEA)^32^ of GGAA microsatellite repeats in peaks that are more accessible in EF^Prx1^ MSCLC than in Wt MSCLC. The barcode indicates peaks with at least eight repeats of the GGAA motif with a variable spacer of 0-3bp. Peaks have been pre-ranked by the DESeq2 test statistic for the comparison of EF^Prx1^ MSCLC and Wt MSCLC. **D** Distribution of log_2_ fold changes (between EF^Prx1^ MSCLC and Wt MSCLC) in peaks separated by the number of repeats of the GGAA microsatellite within these peaks. The plot indicates the kernel density estimate per fold change value, which is similar to a histogram for continuous values.

### IGF-1 exposure assists EF in the stable transformation of limb-derived MSCLC

Although EF^Prx1^ MSCLCs are immortal, they were unable to form colonies when plated at high dilution into soft agar (Figure 3A and B). Consistent with this *in vitro* finding, tumor formation was rarely observed and if, only with long latency, upon transplantation of EF^Prx1^ MSCLC under the skin of SCID (C.B-17/IcrHsd-Prkcdscid) mice (Figure 3C). As the peak incidence of human EwS occurs during adolescence, in which several growth-promoting hormones, including IGF-1 are highly enriched in the bone microenvironment ^22^, we hypothesized that embryonal limb-derived EF^Prx1^ MSCLC may require additional stimuli from the endocrine milieu of the pubertal bone niche for full tumorigenic transformation. We focused on IGF-1 for its proposed essential role in sarcomagenesis. Since there is functional crosstalk between IGF-1R and Insulin signaling in EwS ^16^, we also considered a potential role for Insulin in the transformation process of mouse MSCLC. To test our hypothesis, we plated EF^Prx1^ MSCLC at a density of 1000 cells/3.5 cm^2^ dish in soft-agar containing 10% fetal calf serum (FCS) in presence or absence of supplemented human IGF-1, Insulin, or a combination of both at concentrations of 500ng/ml and 100ng/ml, respectively, reflecting their peak serum levels during human puberty ^33,34^. Hormone supplementation was renewed twice a week. After four weeks of incubation, anchorage-independent colonies became clearly visible in hormone-treated but rarely in control-treated plates of EF^Prx1^ MSCLC (Figure 3A). The number of soft-agar colonies obtained after hormone treatment was similar for EF^Prx1^ MSCLC from both mice (#1 and #2) and about 20-60 times higher than in untreated cultures and did not significantly differ between IGF-1, Insulin, and combination treatments. No colonies were obtained for identically treated Wt MSCLC (data not shown). Since anchorage-independent growth in soft agar is commonly considered a surrogate for malignant transformation, these results suggested that hormone treatment had activated an EF-initiated transformation program. We, therefore, refer to cells from these colonies as “hormone activated” (ha-) EF^Prx1^ MSCLC (Figure 3A).

**Fig. 3.**
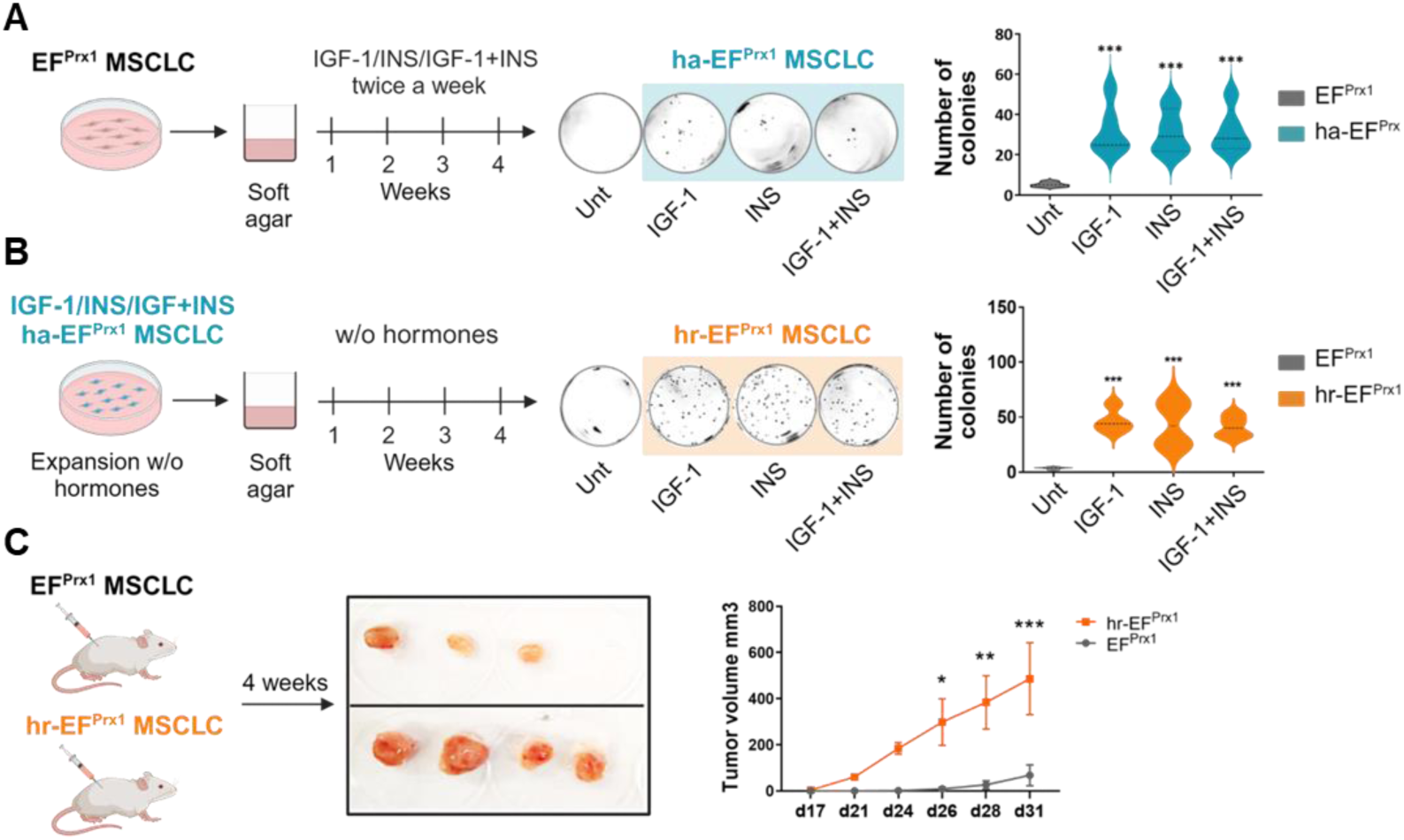
IGF-1 reprogramming of EF^Prx1^ MSCLC to full malignant transformation. **A** Approach assessing *in vitro* anchorage-independent growth of limb-derived MSCLC from EF mutant mice as a surrogate of malignant transformation. EF^Prx1^ MSCLCs were grown in soft agar in the absence or presence of IGF-1 (500ng/ml) and INS (100ng/ml) either alone or in combination according to the indicated scheme. Representative examples of soft-agar plates illustrating the appearance of transformed colonies after four weeks of incubation are shown. Cells derived from these colonies are referred to as hormone-activated (ha-) EF^Prx1^ MSCLC. Colonies > 0.5 mm were counted using ImageJ software. Results are presented as the mean ± SD of triplicate samples (hr-EF^Prx1^ MSCLC#2) from representative data of three independent experiments. (*** P < 0.0005). Unt: untreated (absence of IGF-1/Ins). **B** Same as in **A** but using ha-EF^Prx1^ cells as starting material and culturing cells exclusively in the absence of hormone supplementation. Cells from colonies arising under these conditions are referred to as hormone-reprogrammed (hr-) EF^Prx1^ MSCLC. **C** Tumor formation upon subcutaneous injection of parental EF^Prx1^ #2 and derived hr-EF^Prx1^ MSCLC in SCID mice (4 mice per group). The scheme, representative pictures, and mean tumor size increase over 31 days are shown.

To test if the transformed phenotype required continuous hormone supplementation, we picked ha-EF^Prx1^ MSCLC colonies and expanded them in 10% FCS-containing growth medium for five days in the absence of IGF-1 and Insulin supplementation before re-plating them in hormone-free soft-agar for another four weeks. Strikingly, ha-EF^Prx1^ cells retained and even slightly increased their ability for anchorage-independent growth under hormone-free conditions, while this was not the case for control-treated cells (Figure 3B). We, therefore, refer to stably transformed cells isolated from colonies grown from ha-EF^Prx1^ MSCLC in the absence of further hormone treatment as “hormone reprogrammed” (hr-) EF^Prx1^ MSCLC. Notably, the transformed phenotype of hr-EF^Prx1^ MSCLCs remained dependent on continuous EF expression, as knockdown of the fusion protein drastically reduced their ability to grow in soft agar (Figure S4A). Intriguingly, in contrast to parental EF^Prx1^ MSCLC, hr-EF^Prx1^ MSCLC gave rise to aggressively growing tumors upon xenotransplantation in SCID mice providing evidence for a complete and irreversibly transformed phenotype (Figure 3C).

Wild-type MSCLC, parental non-transformed EF^Prx1^ cells, and their transformed ha-and hr-derivatives were routinely propagated in 10% FCS containing growth medium. FCS comprises multiple hormonal components at low concentrations, with IGF-1 amounts varying around 72ng/ml ^35^. Thus, the final concentration of IGF-1 in the basal growth medium containing 10% FCS is estimated to be about 70 times lower than the concentration (500ng/ml) used for transformative reprogramming of EF^Prx1^ MSCLC. We, therefore, tested the functional status of the IGF-1 signaling pathway in hr-EF^Prx1^ cells. When expanded in 10% FCS containing growth medium, we observed tyrosine phosphorylation of the IGF-1R and serine phosphorylation of its downstream signaling effector AKT, which could be inhibited by the small molecule IGF-1R specific inhibitor NVP-AEW541 ^36^. Under serum starvation, the signal was strongly reduced, consistent with paracrine IGF-1R activation by the minute amounts of hormone contained in complete medium (Figure S4B). Human EwS cell lines are exquisitely sensitive to NVP-AEW541 at IC50 values around 200nM ^37^. In contrast, hr-EF^Prx1^ cells were largely resistant to the IGF-1R inhibitor treatment in adherent (IC50 = 9.3 µM) and spheroid (IC50 =3.7 µM) growth conditions (Figure S4C). However, colony formation in soft agar was greatly reduced at NVP-AEW541 concentrations as low as 0.625µM (Figure S4D). Together, these results suggest that transient exposure to high concentrations of IGF-1 is needed for reprogramming EF^Prx1^ MSCLC to full and stable malignant transformation, while low concentrations are still required for sustained clonogenic growth in soft agar.

### IGF-1 reprogramming enforces an EF transcriptional signature

RNA sequencing was performed for parental EF^Prx1^, ha-EF^Prx1^, and hr-EF^Prx1^ MSCLC to analyze the transcriptomic changes associated with IGF-1 activation and reprogramming. 1,328 and 2,126 (of 55,471) transcripts were found to be up-and downregulated (DESeq2; P_adj_ <= 0.05, |log_2_FC| >= log_2_(2)^30^) in ha-EF^Prx1^ versus parental cells, respectively (Table S6, Figure 4A). Similarly, the expression of 1,079 and 1,732 genes was increased and decreased in hr-EF^Prx1^ MSCLC in comparison to untreated EF^Prx1^ MSCLC, respectively (Table S7, Figure 4A), roughly 42% (up) and 58% (down) of which overlapping with those in ha-EF^Prx1^ MSCLC (Figure 4B). Strikingly, both shared up-and downregulated genes were significantly enriched in human EwS-derived EF target gene sets, including E2F and FOXM1 targets and genes associated with epithelial-mesenchymal transition and extracellular matrix (Figure S5A,B, Table S8). However, the chromatin accessibility of only few genomic regions was affected in hr-EF^Prx1^ MSCLC compared to parental EF^Prx1^ MSCLC (n = 557) (Table S4), and there was no further enrichment in GGAA microsatellites in open chromatin regions, as tested by ATAC-seq (fGSEA; FDR = 0.083)^32^ (Figure 4C,D). This result suggested that IGF-1 reprogramming did not significantly affect the neomorphic activity of EF on GGAA repeat regions but relied on different transcription regulatory mechanisms.

**Fig. 4.**
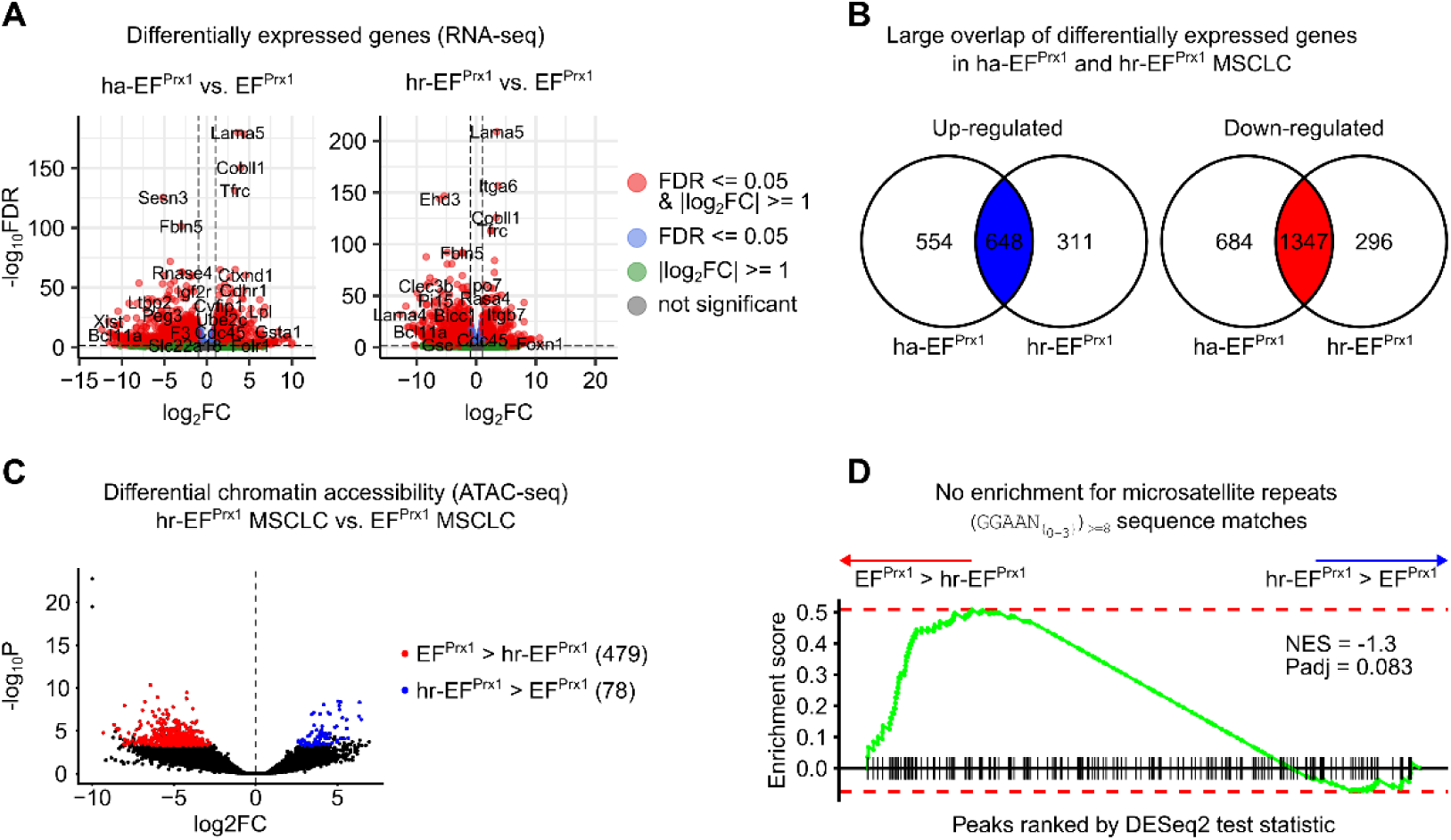
Transcriptional and chromatin changes associated with IGF-1 activation and reprogramming. **A** Volcano plot of differentially expressed genes between EF^Prx1^ MSCLC and ha-EF^Prx1^ (left), respectively hr-EF^Prx1^ (right) MSCLC identified by DEseq2. The X-axis indicates the logarithms of the fold changes of individual genes. The Y-axis indicates the negative logarithm of their P-value to base 10 (DESeq2 ^30^; P_adj_<0.05, |log_2_FC|>log_2_(2); n(EF^Prx1^) = 2, n(ha-EF^Prx1^) = 2, n(hr-EF^Prx1^) = 4). See Tables S6, S7. **B** Venn diagrams showing the overlap of differentially expressed genes between ha-EF^Prx1^ and hr-EF^Prx1^ MSCLC for up-and down-regulated genes in comparison to EF^Prx1^ MSCLC (from **A**). **C** Volcano plot of differentially accessible chromatin regions (from ATAC-seq) between hr-EF^Prx1^ MSCLC and EF^Prx1^ MSCLC. Each point represents an ATAC-seq peak, and significantly up-and down-regulated regions are highlighted in blue and red, respectively (DESeq2^30^; P_adj_<0.05, |log_2_FC|>log_2_(1.5); n(hr-EF^Prx1^) = 2, n(EF^Prx1^) = 4). See Table S5. **D** Fast gene set enrichment analysis (fGSEA ^32^) of GGAA microsatellite repeats in peaks that are more accessible in hr-EF^Prx1^ MSCLC compared to EF^Prx1^ MSCLC. The barcode indicates peaks with at least eight repeats of the GGAA sequences with a variable spacer of 0-3bp. Peaks were pre-ranked by the DESeq2 test statistic for the comparison of hr-EF^Prx1^ MSCLC and EF^Prx1^ MSCLC.

### IGF-1 treatment-related chromatin accessibility changes identify modular mechanisms of EF^Prx1^ MSCLC reprogramming

To understand the transforming mechanisms by which transient IGF-1 exposure permanently reprograms EF^Prx1^ MSCLC, we performed a comparative cluster analysis of open chromatin regions in untreated and IGF-1 treated Wt MSCLC, parental and hr-EF^Prx1^ MSCLC #1 and #2, and one tumor obtained after transplantation of hr-EF^Prx1^ #1 (transplant-hr-EF^Prx1^ MSCLC). Differentially accessible chromatin regions were classified in five distinct clusters designated modules M1 to M5 (Figure 5A; Table S4).

**Fig. 5.**
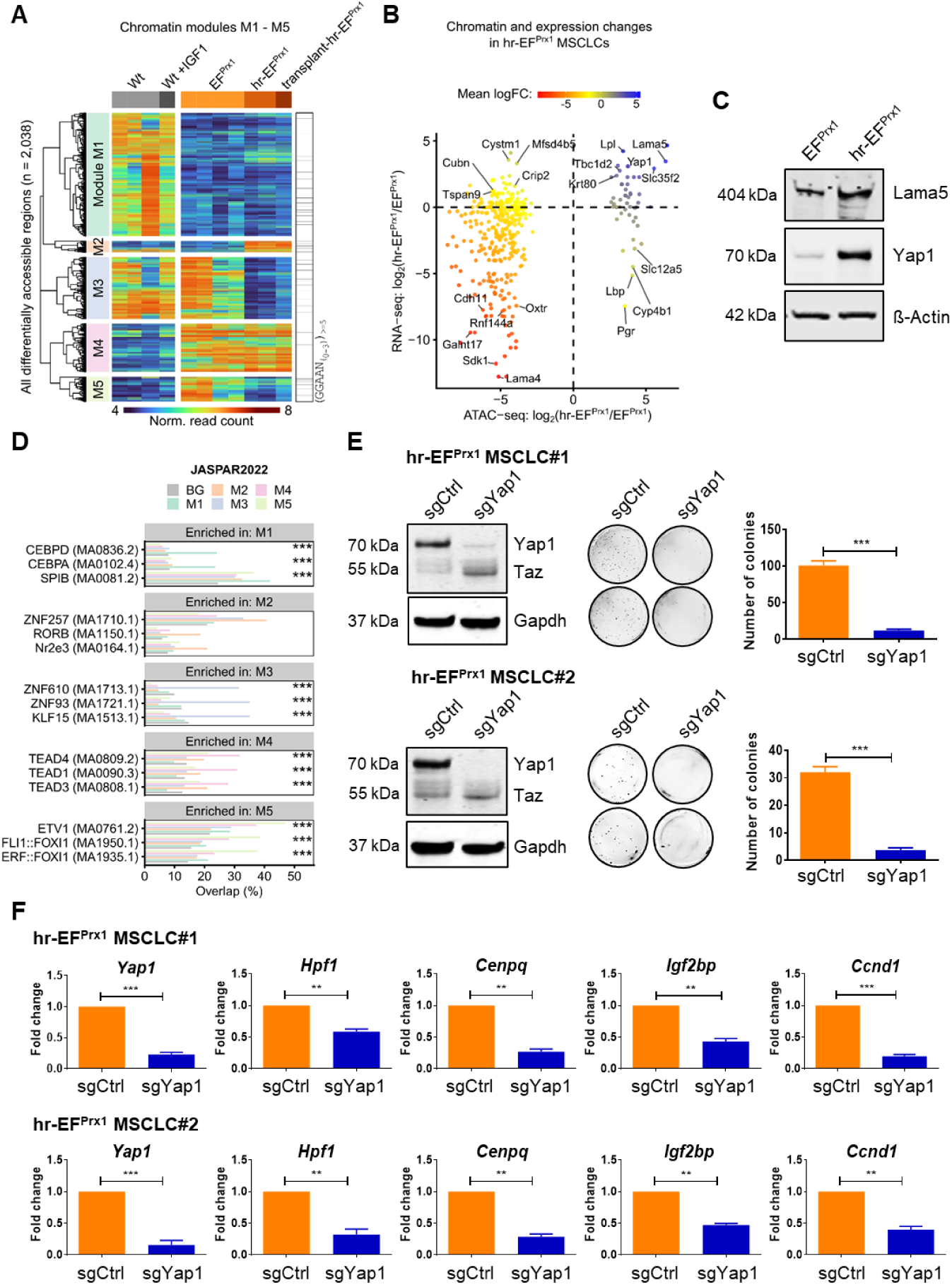
Comparative analysis of differentially accessible chromatin modules in MSCLC. **A** Heatmap showing row-scaled normalized read counts for all peaks that were considered differentially accessible in at least one comparison (EF^Prx1^ MSCLC /Wt MSCLC, hr-EF^Prx1^ MSCLC / EF^Prx1^ MSCLC, or MSCLC+IGF-1/Wt MSCLC; n_total_ = 2, 038). Peaks are grouped into five “modules” (M1-M5). GGAA motif density in differentially open regions is indicated to the right of the heat map. See Table S4. **B** Scatterplot comparing changes in the chromatin accessibility of ATAC-seq peaks (x-axis) with the corresponding changes in gene expression of the nearest genes (y-axis). One point is indicated for each combination of differentially accessible peak (from A) and differentially expressed gene. Color indicates the mean of both fold changes. **C** The protein levels of YAP1, LAMA5, and ß-Actin were detected by Western blot for Wt MSCLC, EF^Prx1^ MSCLC and hr-EF^Prx1^ MSCLC. **D** Bar plots showing DNA sequence motifs (mouse and human TF motifs from JASPAR 2022 ^44^ overrepresented in peaks belonging to each of the five modules from panel A. Each plot panel lists the top 3 motifs per module and each bar indicates the percentage of peaks with at least one match to the given motif. Enrichment was calculated using Fisher‘s exact test (one-sided). *, Padj < 0.05; **, Padj <= 0.01; ***, Padj <= 0.005. See Table S9. **E** Western blot analysis of YAP1 levels in hr-EF^Prx1^ MSCLC#2 cells upon knockout of *Yap1* using three CRISPR sgRNAs (sg-Yap1). Middle panel shows representative soft agar assay for hr-EF^Prx1^ cells transduced with sg-Ctrl vs sg-Yap1. The number of cell colonies was counted on days 21 after plating (right panel). Data is presented as mean ± SE (n = 3), ***p < 0.001. Statistics were calculated by one-sided, paired Student’s t-test. **F** quantitative analysis by RT-qPCR of relative mRNA expression levels of Yap1, Igf2bp1, Hpf1, Cenpq and Ccnd1 after knockout of Yap1. Data is presented as mean ± SE (n = 3), ***P<0.001, **P<0.003. Statistics were calculated by one-sided, paired Student’s t-test.

Cluster M1 comprises genomic regions that are closed in both EF^Prx1^ MSCLC and hr-EF^Prx1^ MSCLC but open in Wt MSCLC irrespective of IGF-1 treatment. Over-represented genes enriched in the vicinity of these regions are involved in inflammatory responses and responses to cytokine stimuli (Figure S6A, Table S9). Vice versa, cluster M4 contains chromatin regions that are closed in Wt MSCLC but open in parental and hr-EF^Prx1^ MSCLC. Top enriched genes in the vicinity of these regions annotate to negative regulation of catabolic processes. In contrast, modules M2 and M3 comprise of genomic regions that are exclusively open (M2) or closed (M3) in hr-EF^Prx1^ MSCLC as a consequence of previous IGF-1 exposure. While there was no preferential enrichment of biological functions for nearby genes of cluster M3, top over-represented M2 genes annotate to steroid hormone signaling. Taking a closer look, we observed that *Yap1* and *Lama5* were the most significant accessible genes among others including EwS hallmark genes *Prkcb* ^38^, *Bcl11b* ^39,40^, and *Sox2* ^41–43^ (Figure S6B). This is consistent with our RNA-seq findings, where there was an enrichment of Hippo pathway genes and specifically a significant up-regulation of *Yap1* and *Lama5* on transcriptional level upon IGF-1 reprogramming, which was corroborated on the protein level by immunoblotting (Figure 5B, 5C). Finally, M5 comprises chromatin regions that are not accessible in Wt MSCLC, open in EF^Prx1^ MSCLC, but which had reversed to a closed state upon IGF-1 reprogramming. Top enriched nearby genes associated with this cluster annotated to protein glycosylation (Figure S6A).

To better understand the gene regulatory mechanisms behind the individual modules, we identified transcription factor DNA binding motifs (from the JASPAR2022 database ^44^) that were enriched in the differentially accessible chromatin clusters (Figure 5D, Table S9). Peaks specifically lost upon EF expression independent of IGF-1 (M1) were enriched in CCAAT box binding factor motifs, while top enriched motifs in those lost upon IGF-1 reprogramming in EF^Prx1^ MSCLC are for KLF15, ZNF610, and ZNF93 in M3, and for PBX2, ETV1, and MEIS2 in M5. The EF and IGF-1 reprogramming-specific module M2 was found enriched in binding motifs for the retinoid acid receptor-related orphan nuclear receptor family, of which we found RorC to be expressed in hr-EF^Prx1^ MSCLC (Figure 5D, Figure S6C). However, this enrichment was not statistically significant. Instead, we observed the highest incidence of GGAA microsatellites in M2. In contrast and unexpectedly, the lowest frequency in GGAA repeats was present in M4 regions, which open up as a consequence of EF expression apparently unaffected by IGF-1 reprogramming (Figure 5A). Interestingly, the top and highly significantly enriched binding motif in M4 is that for Tead transcription factors, which are the downstream nuclear effectors of the Hippo signaling pathway (Figure 5D).

### Sequential EWS::FLI1 and IGF-1 signaling cooperate in the stepwise activation of TEAD target genes

As Tead transcription factor activity requires co-activation by Yap1, which we found to be induced as an M2 component exclusively upon IGF-1 reprogramming in EF expressing MSCLC, we wondered if a fraction of M4 associated genes may become activated in a stepwise manner by the sequential functional activity of the oncogenic fusion protein and IGF-1 signaling. Hence, we screened M4 genes for those that showed highly significant TEAD binding motif accessibility in hr-EF^Prx1^ MSCLC compared to EF^Prx1^ MSCLC. Among them we identified several genes with important functional roles in human EwS biology. We focused on *Igf2bp1,* a top EwS specific dependency in DepMap ^45^, *Hpf1,* an essential co-factor of PARP1/2 which are highly expressed and therapeutic targets in EwS ^46^*, Cenpq,* a gene involved in *FoxM1/Plk-1/Cenp-A* pathway of kinetochor assembly in mitotic progression ^47,48^, and the cell cycle regulator *Ccnd1,* which is overexpressed in EwS ^49^. Indeed, *Yap1* knock-out in hr-EF^Prx1^ MSCLC resulted in downregulation of all these M4 genes consistent with their expression being dependent on *Yap1* activation in M2 (Figure 5E). Together, these results suggest that malignant transformation of embryonal limb-derived MSCLCs in our EF mutant mouse model involved a modular mechanism, in which EF expression opened and primed essential TEAD target genes for subsequent activation by IGF-1-induced Yap1.

To test, if Yap1/Tead interaction was required to maintain the transformed phenotype of hr-EF^Prx1^ MSCLC, we performed soft-agar colony assays in absence and presence of the small molecule TEAD-palmitoylation pocket binder K-975, which was previously shown to inhibit TEAD complex formation with its co-activator YAP1 in mesothelioma ^50^. Colony formation of hr-EF^Prx1^ MSCLC was found to be drastically reduced already at nanomolar drug levels (Figure 6A).

**Fig. 6.**
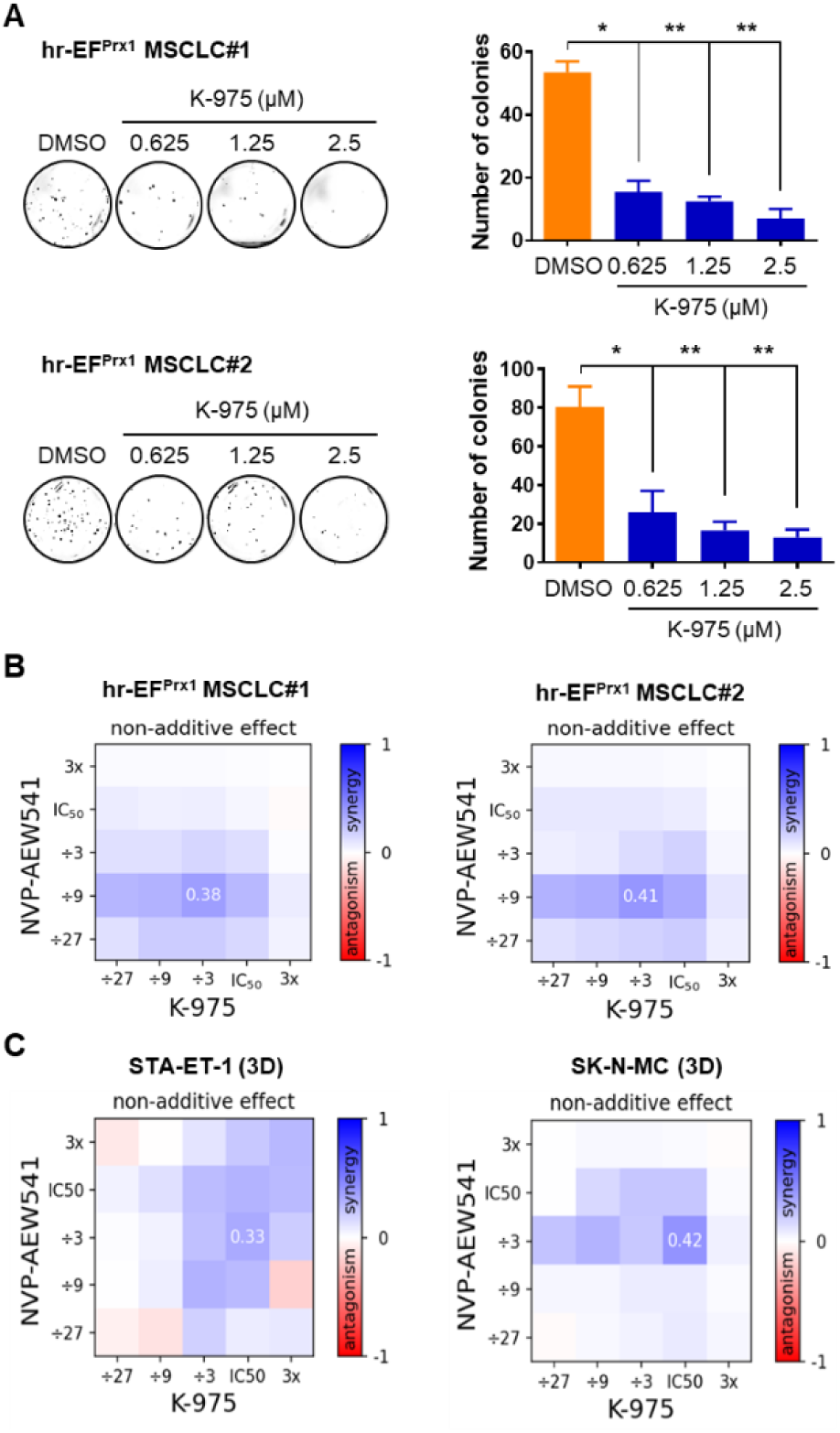
YAP/TEAD blockade reduces colony forming ability and synergizes with IGF-1R inhibition in hr-EF^Prx1^ MSCLC and human EwS. **A** The effects of K-975 on soft agar colony formation of hr-EF^Prx1^ MSCLC upon 12 days incubation; Soft agar assay shows a concentration dependent decrease in the number of hr-EF^Prx1^ MSCLC colonies. Colonies > 0.5 mm were counted using ImageJ software. Data are expressed as mean+/- SD from three experiments, two-way Anova was used to determine statistical significance. **P<0.01, *P<0.05. **B** Overview of synergy scores of NVP-AEW541 and K-975 drug combinations across hr-EF^Prx1^ MSCLC#1 and #2 in monolayer culture conditions. **C** Overview of synergy scores of NVP-AEW541 and K-975 drug combinations for STA-ET-1 and SK-N-MC human EwS cell lines in spheroid culture conditions. The heat map shows Bliss excess across the hr-EF^Prx1^ MSCLC (additive if = 0 and non-additive if ≠ 0; within non-additive cases, it’s synergistic if > 0, or antagonistic if < 0).

Since we had shown that colony formation of established hr-EF^Prx1^ cells remained sensitive to IGF-1R blockade while anchorage-dependent growth was largely resistant to NVP-AEW541 treatment (Figure S4C & D), we tested if Yap/Tead inhibition might increase the sensitivity to IGF-1R blockade. Indeed, combination treatment with K-975 and NVP-AEW541 resulted in synergistic growth inhibition of hr-EF^Prx1^ cells (Figure 6B). Finally, we tested the drug combination in human EwS cell lines TC32 and A673. Results confirm that combined inhibition of IGF-1R and YAP1/TEAD is more effective against EwS than IGF-1R blockade alone (Figure 6C).

### Yap1 amplification supports IGF-1 reprogramming of EF^Prx1^ MSCLC

As preservation of the transformed phenotype of hr-EF^Prx1^ MSCLC required sustained Yap1 activation which we found to be independent of continuous stimulation by high IGF-1, we explored genetic mechanisms enforcing Yap1 expression. Low-coverage whole genome sequencing (lcWGS) of bulk parental EF^Prx1^ cell pools identified subtle copy number gains (less than two-fold) at two regions of chromosome 9q, including the Yap1 locus in the pool derived from one of the two EF^Prx1^ founder mice (#2) but not in the other (Figure S7A). This suggested the presence of a *Yap1* amplified subclone in at least one of the parental cell lines. Analyzing the derivative IGF-1 reprogrammed cell pools, we identified a two-to four-fold copy number increase at the *Yap1* locus for hr-EF^Prx1^#2 cells, and an approximately 8-fold amplification of the same regions in hr-EF^Prx1^ #1 MSCLC (Figure S7B). These results suggest variable expansion of a pre-existing *Yap1* amplified cell clone during IGF-1 reprogramming. To exclude IGF-1 mediated selection of a small Yap1 amplified subclone in parental EF^Prx1^ MSCLC as the origin of stably transformed Yap1 overexpressing cells, we subjected bulk parental EF^Prx1^ MSCLC to minimal dilution and raised single-cell clones. As shown in Figure S7C, none of three tested clones showed chromosome 9 copy number gains consistent with absence of Yap1 protein expression (Figure S7D). However, in each case, IGF-1 reprogramming led to the stable induction of Yap1 and to a lesser extent Taz protein expression (Figure S7D). In contrast to the bulk hr-EF^Prx^ MSCLC populations no amplification of the *Yap1* locus was observed in any of the IGF-1 reprogrammed single-cell clones (Figure S7B), although all of them showed a significant increase in soft agar colony formation similar to the bulk population (Figure S7E). Together, these results suggest that stable induction of Yap1 in response to IGF-1 reprogramming occurs independently of numerical chromosome 9 alterations but may be further supported by *Yap1* copy number gains.

## Discussion

Based on the undifferentiated mesenchymal phenotype, EwS is considered an embryonal malignancy. However, clinical onset of the disease occurs mainly during the second decade of life. While available evidence clearly identifies expression of an EWS::ETS fusion protein as being causal for EwS pathogenesis, developmental timing of this pathognomonic rearrangement is unknown and may occur long before symptomatic disease. It is possible that tumor initiation by the *EWSR1::ETS* gene rearrangement may long precede a tumorigenesis-promoting event. Although p53 mutation was identified as a genetic aberration greatly increasing EF-driven sarcomagenesis in mice and fish ^51,52^, it is rare in EwS patients (<10%) and no other single recurrent genetic or epigenetic aberration has been associated with disease progression in humans, so far.

Here, we tested the hypothesis that tumor initiation by a mutational mechanism happened during embryogenesis leading to differentiation arrest and immortalization of a persisting mesenchymal osteochondrogenic precursor, while tumor promotion occurred by a non-genetic, humoral mechanism in the pubertal bone niche. Such a mechanism is difficult to model *in vivo* as embryonal mesenchymal tissue expression of EF resulted in perinatal lethality. We therefore used explanted embryonal limb-derived MSCLC from EF-mutant mice and subjected them to hormone treatment at a concentration that reflects serum levels during puberty, before assaying them for *in vitro* phenotypic signs of malignant transformation and *in vivo* tumorigenic potential upon transplantation into immune-compromised mice. This model allowed us to identify IGF-1/ Insulin signaling as a tumor promoting mechanism for embryonal EF-expressing MSCLC. Although EF^Prx1^ MSCLC were routinely cultured in presence of 10% FCS, the low IGF-1 concentrations contained in the growth medium did not suffice to promote cellular transformation. In contrast, transient (5 weeks) exposure to high IGF-1 concentrations stably transformed EF^Prx1^ MSCLC to tumorigenicity. The choice of the transforming IGF-1 concentration (500ng/ml) oriented along peak serum levels which are controlled by growth hormone during human puberty ^34^. The actual levels of IGF-1 in the human bone niche, which is produced by various tissues during this developmental period, are not known. During rapid pubertal growth spurt, which among primates is unique to humans ^53^, mechanical load induces IGF-1 production from osteocytes and chondrocytes ^27^, potentially further increasing its levels in the microenvironment of EF-induced EwS precursor cells. Again, this condition is difficult to model in mice, as IGF-1 knockout is perinatal lethal to 95% of offspring ^54^. Even if the IGF-1 concentrations used in our experimental *in vitro* study may not fully reflect the actual in situ levels in the pubertal bone niche, they were sufficient to reproducibly transform EF^Prx1^ MSCLC from two independent founder mice.

Before, it has been doubted that EwS-like tumorigenesis is possible in mice mainly due to differences in GGAA microsatellite landscapes between mice and humans ^2^. Although on average, mice contain more and longer GGAA repeat regions than humans, their number and genomic location relative to essential EwS genes vary significantly. Yet, it has been recently shown that transgenic EF alone can induce EwS-like tumors in the evolutionary much more distant zebrafish ^55^. In our mouse model, we find EF-driven GGAA microsatellite enrichment in open chromatin regions associated with a large number of genes partially overlapping with the human EwS signature. Among them, we find activation of essential GGAA microsatellite-driven EwS hallmark genes *Prkcb* ^38^, the BAF-complex component *Bcl11b* ^39,40^, and the stemness factor *Sox2* ^41^ exclusively upon IGF-1-assisted transformation of EF^Prx1^ MSCLC.

As another potential reason for previous failure to induce EwS tumorigenesis in mice, the well documented toxicity of the EF fusion protein to many cell types was discussed ^2^. Consistent with this supposition, we find downregulation of anti-apoptotic Bcl2 family members and increased apoptosis in EF^Prx1^ MSCLC which, however, did not interfere with their immortalization. Upon IGF-1 reprogramming, we observed decreased apoptosis and induction of anti-apoptotic Mcl-1, which is also found in human EwS providing a potential therapeutic target ^56^.

While transcriptomic, epigenomic and genomic analyses were sufficient to obtain mechanistic insights into the downstream transformation process leading to suppression of developmental programs (M3) and remodeling of stemness genes involved in self-renewal (M2, M4; e.g. Sox2, Klf-5, Ccnd1, Bcl11b, etc.), the actual mechanism of stable transcriptional reprogramming by IGF-1/Insulin remained largely elusive.

Canonical IGF-1 signaling involves activation of the PI3K/AKT-mTOR and the RAS/MAPK-ERK pathways to induce cell growth and survival ^57^. However, in addition to certain HDACs becoming phosphorylated by the PI3K/AKT axis leading to their exclusion from the nucleus more recent evidence identified a direct epigenetic role of nuclear translocated IGF-1R directly binding to and remodeling chromatin via histone H3 phosphorylation and Brg1 recruitment ^58,59^. Whether a similar mechanism might have been involved in the establishment of chromatin accessibility changes specifically found in clusters M2 and M3 of IGF-1 reprogrammed EF^Prx1^ MSCLC remains to be established. In fact, only a small number of chromatin regions got opened in these cells in response to transient high IGF-1 exposure (M2). Here, binding motifs for retinoic acid receptor-like orphan receptors (RORA/B/C) were most prevalent. We found only RORC to be expressed in EF^Prx1^ MSCLC independently of EF and IGF-1, making it unlikely that it was responsible for the observed IGF-1 induced chromatin accessibility changes. However, a recent ChIP-seq study revealed a strong overlap between RORA/C and glucocorticoid receptor (GR) chromatin binding at some 5000 common enhancers and promoters of genes involved in lipid, fatty acid, and amino acid metabolism, the circadian clock and PPARα activity in the mouse liver ^60^. In line, functional annotation of M2 genes identified steroid hormone signaling and lipid metabolism as top enriched terms in this cluster. Intriguingly, EF acts as a transcriptional co-activator for GR in EwS ^61^. GR activation may have also contributed to the observed marked increase in Yap1 expression in EF^Prx1^ MSCLC, similar to what was previously demonstrated for breast cancer in humans, where GR antagonists reduced YAP levels and inhibited cancer stem cell formation ^62,63^. Consistent with our findings there is evidence for an evolutionary conserved PI3K dependent but AKT largely independent mechanism of Yki/Yap induction by IGF-1/Insulin signaling, which feeds back on IGF-1/Insulin signaling via direct upregulation of the Insulin receptor ^64^. Additionally, more recent studies in liver cancer and diffuse large B-cell lymphoma revealed stabilization and activation of YAP1 nuclear translocation in response to IGF-1R signaling associated with bad prognosis^65^.

Insulin/IGF-1 signaling through PI3K is also required in the activation of Yki/Yap by mechanical and polarity cues ^66^. Mechanical tension is mediated via extracellular matrix (ECM) components communicating with integrins and cell adhesion molecules on the cell surface, leading through activation of Focal adhesion and Src kinases to LATS1/2 dependent YAP1 activation ^63^. The major ECM component involved in stem-cell self-renewal, migration, adhesion and differentiation is Laminin. As demonstrated for human iPSC cells, culturing on Laminin 511-containing substrate prevents chondrogenic differentiation through a mechanism involving nuclear accumulation of unphosphorylated YAP1 ^67^. We find the Laminin 511 component LAMA5 to be highly induced by the combined activity of EF and IGF-1 in EF^Prx1^ MSCLC impaired in osteogenic and adipogenic differentiation potential and unable to complete *in vivo* chondroblastic differentiation. LAMA5-containing laminins interact with integrin receptors containing the β1 subunit ^51^. Enhanced LAMA5 and β1 integrin expression was previously reported to be induced by IGF-1 treatment of cultured corneal epithelial cells ^68^. Endogenously produced LAMA5 promotes self-renewal of human pluripotent stem cells in an autocrine and paracrine manner through E-cadherin and FYN-RhoA-ROCK activation ^69^, and it is tempting to speculate that it fulfills a similar function in hr-EF^Prx1^ MSCLC. Of note, the RhoA GAP Arhgap42, which was recently demonstrated to be involved in ECM signaling under mechanical force ^70^, was also among highly upregulated hr-EF^Prx1^ specific M2 genes in our study. Expression levels of the Laminin 511 receptor α6β1 integrin and of E-cadherin were reported to be controlled among others by IGF-1R signaling resulting in maintenance of pluripotency and self-renewal ^71^. Likewise, the activation of the Laminin receptor of human MSC was shown to regulate OCT4 and SOX2 stem cell factor expression^72^. In our study, chromatin accessibility at the *Sox2* locus was increased in hr-EF^Prx1^ cells together with activation of a further stem cell factor, KLF5 (Figure S8A, Table S5), encoded in chromatin accessibility cluster M4 and known to be stabilized by Yap1 protein ^73^. In fact, knockout of *Yap1* in hr-EF^Prx1^ cells resulted in a decrease in KLF5 protein levels, which could be rescued upon pharmacological proteasome inhibition (Figure S8A). YAP1 appeared also involved in the observed upregulation of FOXM1 and its transcriptional program as FoxM1 RNA and protein expression was greatly reduced upon Yap1/Tead inhibition by sgRNA or K-975 (Figure S8B). FOXM1 is a known mediator of EF dependent cell cycle regulation in EwS ^74^. Together, these results suggest that IGF-1 reprogramming of EF^Prx1^ MSCLC enforces a stem cell program as the basis of malignant transformation.

The major finding of our study was that of a modular mechanism of stepwise malignant transformation by EF and IGF-1/Insulin. In the first step, EF rewired the transcriptome of an embryonal osteochondrogenic precursor resulting in developmental arrest and cellular immortalization. Here, the neomorphic activity of EF to bind and open GGAA-microsatellites may have played an important role which was, however, not sufficient to mediate full transformation. Instead, a large cluster of genes devoid of GGAA repeat regions but enriched in TEAD binding motifs (M4) opened in response to EF expression priming them for binding and activation by the respective transcription factor. However, the actual activation of these genes occurred only upon induction of the co-activator Yap1 in the second reprogramming step following transient stimulation by high IGF-1/Insulin. Although not required for IGF-1-driven full transformation, copy number gain at the *Yap1* gene locus if present in a subclone of parental EF expressing MSCLC cells was positively selected in hr-EF^Prx1^ bulk cell populations. *Yap1* amplification frequently occurs during human and mouse tumorigenesis though it has not yet been reported for EwS despite being highly expressed and associated with poor prognosis in this disease ^75^ ^76–83^.

We hypothesize that Yap1 activation as a rate-limiting tumorigenic step occurs in the bone niche during puberty and further leads to epigenomic reprogramming resulting in suppression of heterogenous developmental genes by an as yet unknown mechanism (M3, M5), and the direct and indirect activation of a number of stemness and dependency genes involved in cell renewal including *Sox2*, *KLf5*, and *Igf2bp1,* as well as of key genes involved in cell cycle (i.e *Ccnd1*), mitotic progression (i.e. *FoxM1, Cenpq*) DNA-repair (i.e.*Hpf1*), and survival (*Mcl-1*). Thus, it appears that the Hippo pathway played a key role in tumor initiation in our model. In contrast in established human EwS tumors, this pathway has been associated with epithelial/mesenchymal-like transition (EMT) and metastasis being activated upon EF modulation. In fact, in response to shRNA-mediated EF downregulation we previously observed activation of TEAD target genes by co-activators MRTFB and the YAP1 ortholog TAZ (WWTR1), while YAP1 was invariably expressed in all tested EwS cell lines consistent with immunohistochemical positivity in a large fraction of primary EwS tumors. Although the current scientific literature on Hippo pathway biology rarely distinguishes between YAP1 and TAZ, there is accumulating evidence that the two orthologous proteins may have different functions and regulate different TEAD target gene sets during normal limb development ^83–85^. We therefore hypothesize that YAP1 may be essential for the onset of disease during puberty, while TAZ may drive metastasis of EwS. We therefore expect both activities to be sensitive to small molecule inhibition of TEAD complex formation with its alternative co-activators YAP1 and TAZ, and accordingly we have already shown that the metastatic potential of EwS is reduced by treatment with the YAP/TAZ/TEAD inhibitor verteporfin ^86^. Here, we show that an allosteric TEAD inhibitor binding specifically to the TEAD palmitoylation pocket, K-975, greatly reduced the anchorage-independent colony forming ability of hr-EF^Prx1^ MSCLC and of EwS cell lines *in vitro*. Several related TEAD inhibitors have recently entered clinical trials for the treatment of high risk solid tumors ^87^. Our study provides a mechanistic rationale to combine YAP1/TEAD targeting compounds with IGF-1R directed therapy to improve outcome of EwS patients. Here, we provide first proof-of-principle *in vitro* evidence for the efficacy of this combination in our model and in EwS cell lines encouraging further pre-clinical development.

## Methods

### Genetically modified mice

Mice harboring a Cre-inducible *EF* knocked into the ubiquitous *Rosa26* locus (*Rosa26loxP-STOPloxP-HA-EF* allele) ^28^ were crossed to a Prx1Cre line ^29^. To determine the age of the embryos, female mice in breeding cages were daily checked for vaginal plugs. If a plug was found, the female was separated at noon and that day was counted as E (embryonic day) 0,5. Embryos were then harvested at indicated time points. Mice were kept under standard conditions at the Decentralized Biomedical Facility of the Medical University of Vienna on a 12 hours light-dark cycle. All animal experiments were approved by the Austrian ministry for science and research (license number: BMWF-66009/0139-C/GT/2007).

### Cell lines

Among EwS cell lines used in this study, STA-ET-1 was established in-house, TC32 was kindly provided by Sue Burchill (University of Leeds, UK), A673 and SK-N-MC were from the American Type Culture Collection (CRL-1598 and HTB-10). Lenti-X293T lenti-viral packaging cells were purchased from Takara Bio (# 632180). Cell lines were cultivated in RPMI1640 (# 61870044, Thermo Fisher Scientific) and DMEM (# 2023-09, Gibco) in the presence of 10% fetal calf serum (#10270106, Thermo Fisher Scientific).

### Preparation and expansion of MSCLC

Mesenchymal stem cell like cells (MSCL) from Prx1Cre founder mice used as controls were isolated by flashing the femur and tibia of P1 day old mice. To obtain the cells from cartilaginous elements of mouse EF^Prx1^ mutants, they were first cleaned from muscles and surrounding tissues, then cut in pieces and left in alpha-DMEM, 10% FCS, 0.1% Pen/Strep and 0.1% glutamine.

### Skeletal staining

Embryos were skinned, eviscerated, fixed in 4% paraformaldehyde (PFA)/PBS at 4°C overnight and incubated in 96% ethanol. Fat was removed by incubating in acetone and embryos were stained with 0,015% Alcian Blue (A5268-10G, Sigma-Aldrich) /0,005% Alizarin Red (a5533-25g, Sigma-Aldrich) in 5% acetic acid, 60% ethanol. Surrounding tissue was cleared with 1% potassium hydroxide and skeletons were stored in glycerol.

### Histology

Tissues were fixed in 4% PFA/PBS overnight at 4°C, dehydrated, embedded in paraffin and cut into 5µm thick slices. Sections were stained with hematoxylin (SLCN6532, Sigma-Aldrich) and eosin (SLCP2819, Sigma-Aldrich) by using standard protocols and analyzed with a Zeiss Axio Imager.Z1 microscope.

### RNA in situ hybridization

Digoxogenin labeled cRNA probes were generated by *in vitro* transcription of 1µg template DNA with the RNA DIG labeling kit (#11175025910, Roche) according to the manufacturer’s protocol. Tissue slides were de-waxed with Shandon Xylene Substitute (#9990505, Thermo Scientific™), re-hydrated, fixed with 4% PFA/PBS and proteins were removed by incubation with Proteinase K (P2308, Sigma-Aldrich). After acetylation with acetic anhydride (#242845, Sigma-Aldrich) the labeled probe was mixed with hybridization buffer (10mM Tris ph 7,5; 500mM NaCl; 1mM EDTA; 0,25% SDS; 10% Dextran sulphate; 1x Denhardts’s (0,02% Ficoll 400; 0,02% Polyvinylpyrolidone; 0,02% BSA); 200µg/ml yeast tRNA; 50% formamide), applied to the slide and hybridization was carried out at 65°C. RNA was digested and slides were washed with decreasing salt concentrations. Specific hybridization was visualized by incubating slides with anti-Digoxigenin-Alkaline phosphatase antibody (1:2000) (#11093274910, Roche) and BM purple AP substrate (#11442074001, Roche). The reaction was stopped with NTMT (100mM NaCl; 100mM Tris pH 9,5; 50mM MgCl2; 0,1% Tween–20), slides were covered with glycergel mounting medium (C0563, Dako).

### Flow cytometry

For identifying mesenchymal markers of bone marrow derived cells, Wt MSCLC and EF^Prx1^ MSCLCs were harvested with Accutase (A1110501, Gibco). A total of 0.5 × 10^6^ cells were washed with PBS then transferred to FACS tubes and centrifuged for 5min at 400g. Cells were incubated with specific individual monoclonal antibodies, conjugated with phycoerythrin (PE), allophycocyanin (APC) and allophycocyanin/cyanine7 (APC/Cyanine7) in CliniMACS-buffer (130-021-201, Miltenyi Biotec GmbH) for 30 min in the dark at room temperature. The following cell surface antigens were assessed: CD90-2-APC (17-0902-82, eBioscience™), CD44-PE (12-0441-82, eBioscience™), CD45-APC (17-0454-82, eBioscience™) and CD19-APC-Cy7 (115501, Biolegend). Mouse isotype-matched IgG served as a negative control. Flow cytometry was performed on a *BD LSRFortessa™ Cell Analyzer* and data were processed with FCS express software 7.

### Adipogenic, osteogenic and chondrogenic differentiation assay

1 × 10^5^ Wt MSCLC and EF^Prx1^MSCLCs were seeded into a six-well plate. For adipocyte differentiation, we used MesenCult medium containing 10% adipogenic stimulatory supplements (#5507, Stem Cell Technologies). At day 21, cells were rinsed with PBS twice and fixed with 4% PFA for 60 min. Cells were then rinsed with distilled water and incubated in 60% isopropanol for 2 min. Finally, cells were covered with Oil Red O solution (O0625, Sigma-Aldrich) for 5 min. Osteogenic differentiation was performed by using the MesenCult osteogenic stimulatory kit (#5504, Stem Cell Technologies). Medium was changed every 2 days, and the cells were analyzed after 21 days of differentiation. After fixation with PFA for 60 min, cells were washed with ddH2O and stained with fresh Alizarin Red S solution and incubated at room temperature in the dark for > 45 minutes followed by 4 washing steps with ddH_2_O. Chondrogenic differentiation was achieved using the MesenCult-ACF Chondrogenic Differentiation Kit (#5455, StemCell Technology). After fixation with PFA for 60 min, cells were rinsed with PBS and stained with 1% Alcian blue solution prepared in 0.1 N HCL for 30 minutes followed by 3 washing steps with water.

### Soft agar assay

For soft agar assays, each well of a twelve-well plate was first covered with an underlayer of 0.6% SeaPlaque GTG agarose (#50111, Lonza,) in growth medium (RPMI1640 supplemented with penicillin/streptomycin and glutamine). Then, cells were seeded at a density of 1 × 10^3^ cells per well in growth medium containing 0.3% agarose. Upon agarose solidification, plates were incubated at 37°C and 5% CO2%. After 4 weeks, plates were imaged, and colonies were counted.

### *In vivo* tumor formation assay

2.5 × 10^6^ cells/ml in PBS were subcutaneously injected into immunodeficient SCID mice (C.B-17/IcrHsd-Prkcdscid; Charles River Laboratories). Tumor formation was examined periodically by palpation. The tumor volume was calculated from tumor size by the formula (diameter × diameter × length/2). The animals were sacrificed, and tumors were surgically removed 31 days after injection. All experiments were performed according to the Austrian guidelines for animal care and protection and were approved by the Austrian ministry for science and research under the license number BMBWF-66.009/0233-V/3b/2019. 2-way anova was used to compare tumors formed by hr-EF^prx1^MSCLCs and EF^prx1^MSCLCs.

### SiRNA knockdown

For silencing of Yap1, cells were transfected with ON-TARGETplus SMARTpool mouse siRNAs against Yap1 (#L-012200-00-0005, Dharmacon/Horizon) using Oligofectamine reagent (#58303, Invitrogen/Thermo Fisher Scientific). As a control, ON-TARGETplus non-targeting pool (D-001810-10-20, Dharmacon/Horizon) was used. After 24h, the transfection procedure was repeated, and cells were collected for qPCR and Western blot.

### Production of viral Cas9 and sgRNA expression vectors

Lenti-X293T cells were transfected with 10 µg of LentiCRISPR /Yap1 sgRNA Crispr/Cas9 all in-one lentivector set mouse (#50584114, abcam), 5 µg of psPAX2 (#12260, Addgene), and 5 µg of pMD2.G (#12259, Addgene) using Lipofectamine 2000 (#11668019, Invitrogen Corporation) according to manufacturer’s instructions. After at least 48 hours of incubation, the virus-containing supernatant was harvested and concentrated using Lenti-X™ Concentrator (#631231, Takara) at 4°C overnight. After centrifugation at 1,500 x g for 45 minutes at 4°C, the pellet was resuspended in one hundredth of the original volume using complete RPMI1640 medium. 10^5^ target cells were seeded per well in a 24-well plate for 24 hours prior to viral infection with 0.5 ml of virus suspension in complete medium in presence of 8 μg/ml Polybrene (TR-1003-G, Millipore Sigma).

### Drug synergy assay

For estimation of synergy we used an established Bliss drugs’ independence model. According to the Bliss model, drugs act independently if the surviving fraction of cells upon simultaneous administration is equal the product of surviving fractions when drugs are given separately. In order to capture complex drug interaction patterns across dose pairs, dose–response matrices were used, as synergy may exist only for specific pairs of treatment doses. Triplicate dose matrices were generated for each drug pair, positioned in different screening plates (ViewPlate-96, White 96-well Microplate with Clear Bottom, #6005181, PerkinElmer) and based on the fold dilutions of the 72h IC_50_ values (determined beforehand for each drug). Cell viability was determined after combinatorial drug treatment using CellTiter-Glo® Luminescent Cell Viability Assay (#G7570, Promega), and the Bliss-predicted inhibition was calculated and compared to the observed values to quantify the interaction between the drugs for each matrix position (additive if = 0 and non-additive if ≠ 0; within non-additive cases, it is synergistic if > 0, or antagonistic if < 0).

### RNA sequencing

#### Library preparation and sequencing

To prepare samples for RNA sequencing, total RNA was isolated using Trizol or RNeasy Mini Kit (#74106, QIAGEN). The library was prepared from these samples by poly(A) enrichment and sequenced with 50 bp single-end read mode on an Illumina HiSeq instrument at the Biomedical Sequencing Facility (BSF) at the CeMM Research Center for Molecular Medicine of the Austrian Academy of Sciences (data for Fig. S3) or the Vienna BioCenter Next Generation Sequencing Core Facility (VBCF-NGS; all other RNA-seq data).

#### Data processing and analysis

Raw sequencing data were processed using the nf-core/rnaseq v3.14.0 pipeline^88^, including alignment to the GRCm38.p6 mouse reference genome and gene-level read quantification using the Ensembl v99 gene annotations. Following initial data processing, all subsequent analyses were performed in R v4.2.2 using Bioconductor packages. The read counts were loaded into DESeq2 ^30^ v1.38.3 or variance-stabilizing transformation and differential analysis (default parameters). Transcripts with an FDR-adjusted P-value <= 0.05 and an absolute log_2_ fold change >= 1 were considered significant. All differential expression analysis results are reported in Tables S2, S6 and S7. For functional enrichment analysis we performed hypergeometric tests using the hypeR package v1.14.0 ^89^ and the following databases from enrichR ^90^: "TF_Perturbations_Followed_by_Expression", “MSigDB_Hallmark_2020”, “ChEA_2022”, and "KEGG_2021_Human". As a background for this analysis, we used all genes identified in our analysis. Terms with an FDR-adjusted P-value <= 0.05 were considered significant. For gene set enrichment analysis, we used fGSEA ^32^ v1.24.0 and the DESeq2 test statistic as a ranking criterion. Gene signatures from Kinsey at al ^91^. and Riggi et al ^92^. were obtained from MSigDB ^93^. All enrichment results are reported in Tables S3 and S8.

### ATAC-Seq

#### Library preparation and sequencing

ATAC-seq was performed following published protocols ^31^. Briefly, 20,000 to 50,000 cells were lysed in a buffer containing digitonin (#16359, New England Biolabs) and Tn5 transposase enzyme (Tagment DNA Enzyme and Buffer Kit, #20034197, Illumina). After incubation at 37°C for 30 minutes, tagmented DNA was purified and enriched. ATAC-seq libraries were prepared for samples of the following experimental conditions: wt MSCLC (n = 5; N.B., one sample failed quality control in the subsequent analysis, see below), wt MSCLC with IGF-1 (n = 1), EF^Prx1^ MSCLC (n = 4), hr-EF^Prx1^ MSCLC (n = 2), transplant-hr-EF^Prx1^ MSCLC (n = 1) for sequencing. Sequencing was done in 50 bp single-end read mode on an Illumina HiSeq instrument at the Biomedical Sequencing Facility (BSF) at the CeMM Research Center of Molecular Medicine of the Austrian Academy of Sciences.

#### Data processing and analysis

Raw sequencing data were processed using pypiper v0.10.0 and PEPATAC v0.8.6 ^94^ with MACS2 v2.1.0 ^95^ for peak calling. Following initial data processing, all subsequent analyses were performed in R v4.2.2 using Bioconductor packages. Only samples with a Non-Redundant Fraction (NRF) >= 0.5, a PCR bottlenecking coefficients PBC1 >= 0.7 and PBC2 >= 3, and an TSS enrichment score >= 10 were considered for further analysis (one wt MSCLC sample failed these criteria; see https://www.encodeproject.org/data-standards/terms/ for a definition of these terms). After merging peaks across all ATAC-seq datasets and removing peaks that overlapped blacklisted regions from ENCODE (https://sites.google.com/site/anshulkundaje/projects/blacklists), we counted for each input dataset the number of reads overlapping the retained peaks with featureCounts (Rsubread v2.12.3) ^96^. The raw read counts were loaded into DESeq2 ^30^ v1.38.3 or variance-stabilizing transformation and differential analysis (using a “batch” as a covariate; other parameters: lfcThreshold=log2(1.5), independentFiltering=TRUE). Peaks with an FDR-adjusted P-value <= 0.05 were considered significant. All ATAC-seq peaks and differential accessibility results are reported in Tables S4 and S5, respectively. For functional enrichment analysis we performed hypergeometric tests using the hypeR package v1.14.0 ^89^ and the following databases from enrichR ^90^: "GO_Biological_Process_2021", "PanglaoDB_Augmented_2021", "TF_Perturbations_Followed_by_Expression", and "TRRUST_Transcription_Factors_2019". As a background for this analysis, we used all genes associated with at least one peak in our analysis (i.e., genes whose transcription start site was within 100kb of a peak). Terms with an FDR-adjusted P-value <= 0.005, a log2 odds ratio >= log2(2), and with >= 10 genes in the overlap were considered significant. For motif analysis, we used motifmatchr v1.20.0 ^97^ to scan all ATAC-seq peaks for matches to known motifs from the JASPAR2022 ^97^ database (R package v0.99.8). We then used base R functions to calculate Fisher’s exact test (one-sided), considering motif hits with an FDR-adjusted P-value <= 0.005, a log2 odds ratio >= log2(2), and occurring in >= 5% of peaks as significant. To identify GGAA microsatellite repeats, we used a regular expression. We then used fast gene set enrichment analysis (fgsea ^32^ v1.24.0) using the DESeq2 test statistic as a ranking criterion to test for enrichments. All enrichment results are reported in Table S9.

### Low coverage whole genome sequencing (lcWGS)

#### Library preparation and sequencing

lcWGS was performed for Wt MSCLC, EF^Prx1^ MSCLC and hr-EF^Prx1^ MSCLC. Genomic DNA was extracted using the Monarch® Genomic DNA Purification Kit (#T3010S) and sequencing service was utilized running on an Illumina NovaSeq 6000 platform. A minimum of 100 ng of DNA was sent to and sequenced by Eurofins Genomics. This service included preparation of a 450 bp DNA sequencing library using a modified version of the NEBNext Ultra™ II FS DNA Library Prep Kit for Illumina and sequencing on an Illumina NovaSeq 6000 with S4 flowcell, XP workflow and in PE150 mode (Illumina).

#### Data processing and analysis

Raw sequencing data were processed using the nf-core/sarek v3.2.1 ^98^ pipeline and aligned to the Mus musculus mm10 reference genome. Copy number variations were inferred using the CNVKit toolkit v0.9.9 ^99^, integrated into the sarek v3.2.1 pipeline ^98^. Visualization of the results for specific regions of interest was carried out through a custom Python3 script.

## Data and code availability

The RNA-seq and ATAC-seq data generated in this study will be deposited in the Gene Expression Omnibus (GEO). Computer code used for data analysis will be shared via our GitHub organization (https://github.com/cancerbits).

## Acknowledgements

We thank Drs. S. Baker and M. Logan for providing EF knock-in and Prx1-Cre transgenic mice crossed to generate EF^Prx1^ mice.

Funding: This study was supported by European Uniońs Horizon 2020 Innovative Medicines Initiative H2020-lMI2-JTl-201 5-07, grant 116064 (ITCC P4), European Union’s Horizon 2020 research and innovation programme under the Marie Skłodowska-Curie grant agreement 956285 (VAGABOND), the Alexs Lemonade Stand Foundation (ALSF) “Crazy 8” grant 20-17258, and grants from the Austrian Science Fund FWF (P34341-B and P35353-B) to H.K.

## Author contributions

Conceptualization, H.K., and R.M. Data curation, F.H., and M.K. Formal Analysis, R.N., A.B., M.K., and F.H. Funding acquisition, H.K. and R.M. Investigation, R.N., B.S., B.R.S., V.F., M.S., L.K., G.W., V.S., and T.J. Methodology, T.J., B.S., and B.R.S. Project administration, H.K. Resources, T.J., W.M., L.K., and R.M. Software, F.H., and A.B. Supervision, H.K., and R.M. Visualization, R.N., F.H., A.B., and V. F. Writing – original draft, R.N., and H.K. Writing – review & editing, F.H., B.R.S., V.F., R.M.

## Supplementary information

Supplementary Figures 1-8

Supplementary Tables 1-9. Excel file containing additional data too large to fit in a PDF.

**Correspondence** and requests should be addressed to Heinrich Kovar

## Supplementary Figures

**Supplementary Figure 1.**
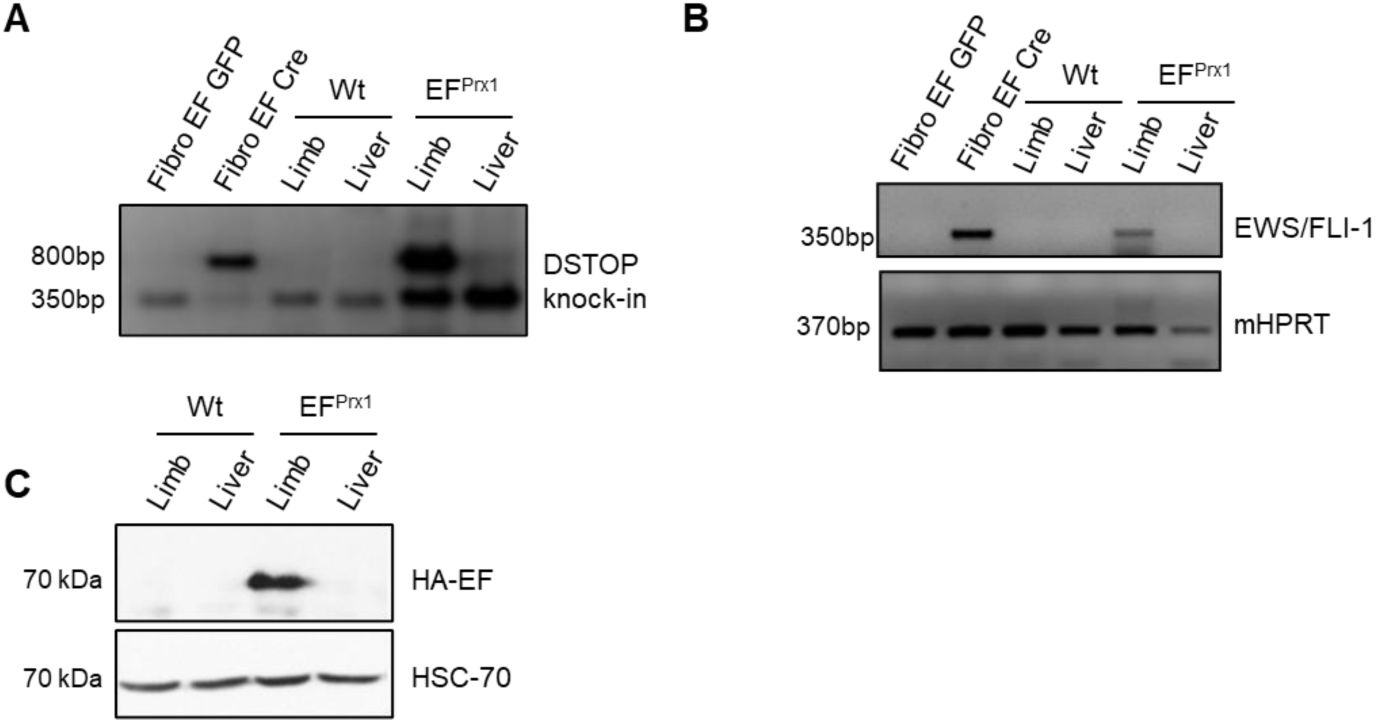
EF expression in the developing limb of EF^Prx1^ mice. **A** Analysis of genomic DNA of wild type (Wt) and EF^Prx1^ mice (E17.5) for deletion of STOP-cassette (DSTOP) with a PCR strategy. **B** Detection of EF expression from reverse transcribed cDNA with EF-specific primers. mHPRT was used as an internal control. Fibro EF GFP or Cre: fibroblasts isolated from EF mice lentivirally transduced with an expression construct encoding GFP or Cre recombinase served as negative and positive controls. **C** The presence of EF protein in EF^Prx1^ limbs was verified by immunoblot analysis using an antibody specific for the HA-tag. HSC-70 was used as a loading control.

**Supplementary Figure 2.**
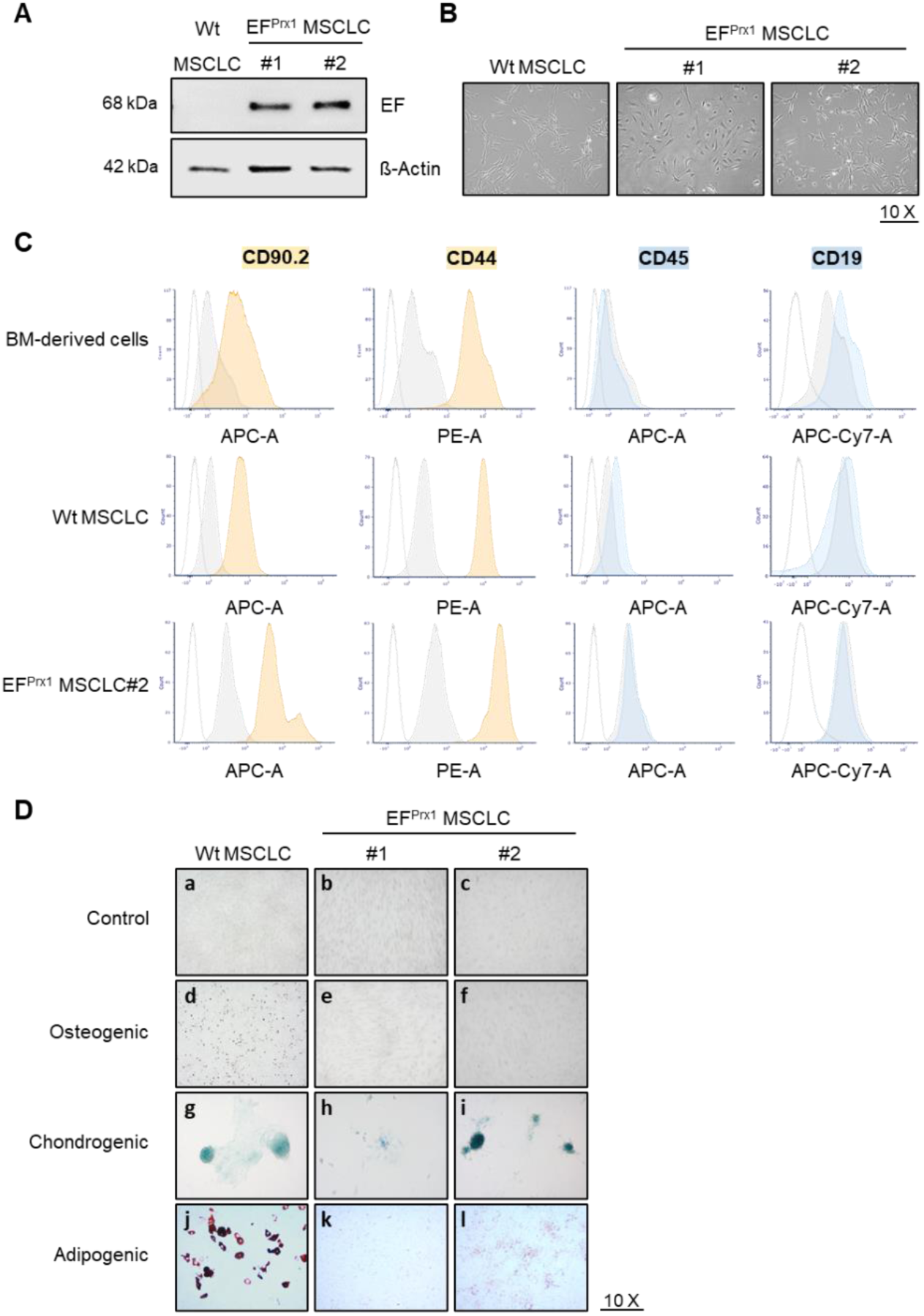
Differentiation potential of EF expressing bone derived MSCLC. EF^Prx1^ MSCLCs stably expressed EF protein and were morphologically and immunophenotypically similar to Wt MSCLC. **A** Presence of EF protein in MSCLC isolated from EF^Prx1^ limb bud (EF^Prx1^ MSCLC) was verified with Western blot analysis using an antibody against the FLI1 C-terminus. MSCLC from the limb of Prx1Cre mice (Wt MSCLCs) served as a negative control. β-Actin was used as a loading control. **B** Morphology of Wt and EF^Prx1^ MSCLC#1 and #2 (magnification 10×). **C** Flow cytometric characterization of Wt MSCLC and EF^Prx1^ MSCLC revealing positivity for mesenchymal markers CD90 and CD44 (orang-filled peaks) and negativity for hematopoietic markers CD45 and CD19 (blue-filled peaks). Grey-filled peaks show staining with isotype controls; empty peaks show negative controls of unstained cells. **D** *In vitro* differentiation potential of MSCLC and EF^Prx1^ MSCLC#1 and #2 upon 21 days incubation with differentiation inducing cocktails for osteogenic, chondrogenic, and adipogenic differentiation. (**a**–**f**) Alizarin red staining, (**g**–**i**) Alcian blue staining, and (**j**–**l**) Oil Red O staining was used to assess osteogenic, chondrogenic, and adipogenic differentiation, respectively (magnification 10×).

**Supplementary Figure 3.**
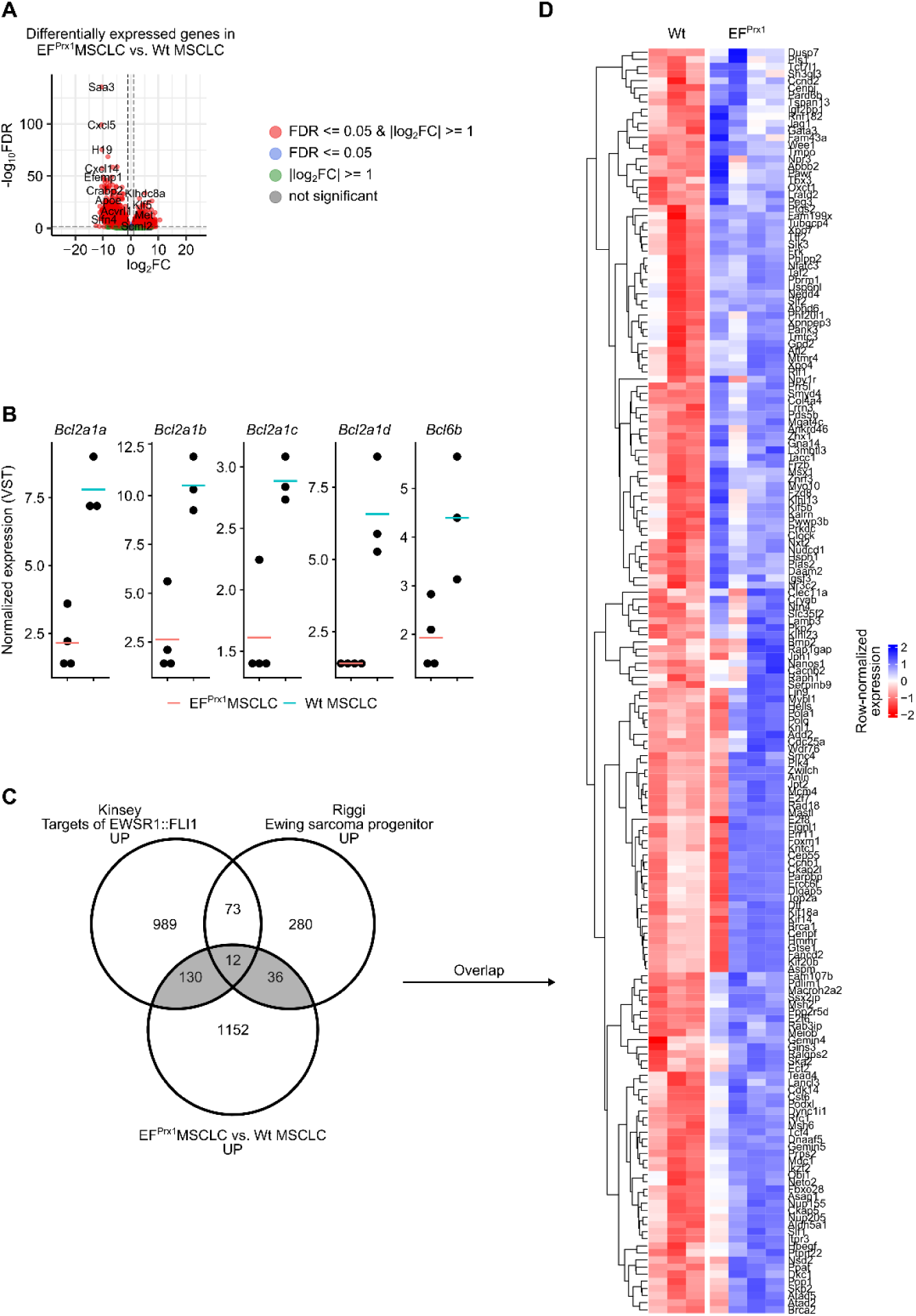
Differentially expressed genes between EF^Prx1^ MSCLC and Wt MSCLC. **A** Volcano plot of differentially expressed genes between EF^Prx1^ MSCLC and Wt MSCLC identified by DEseq2. The X-axis indicates the logarithms of the fold changes of individual genes. The Y-axis indicates the negative logarithm of their P-value to base 10 (DESeq2 ^30^; P_adj_<0.05, |log_2_FC|>log_2_(2); n(EF^Prx1^) = 4, n(Wt) = 3). See Table S2. **B** Normalized RNA-seq counts of transcripts for Bcl2a family members in EF^Prx1^ MSCLC *vs*. Wt MSCLC groups. **C** Venn diagrams showing the overlap between up-regulated genes in EF^Prx1^ MSCLC vs Wt MSCLC and genes modulated by EF upon knockdown in EwS cell lines TC71 and EWS502 ^91^, and genes upregulated in human mesenchymal stem cells engineered to express ectopic EF ^92^. Overlaps between human and mouse gene signatures were determined by matching gene symbols. See Table S3 for results of overlap enrichment analyses. **D** Heatmap showing z-score of DEseq2 normalized counts for candidate EF target in the overlap of gene signatures shown in **C**.

**Supplementary Figure 4.**
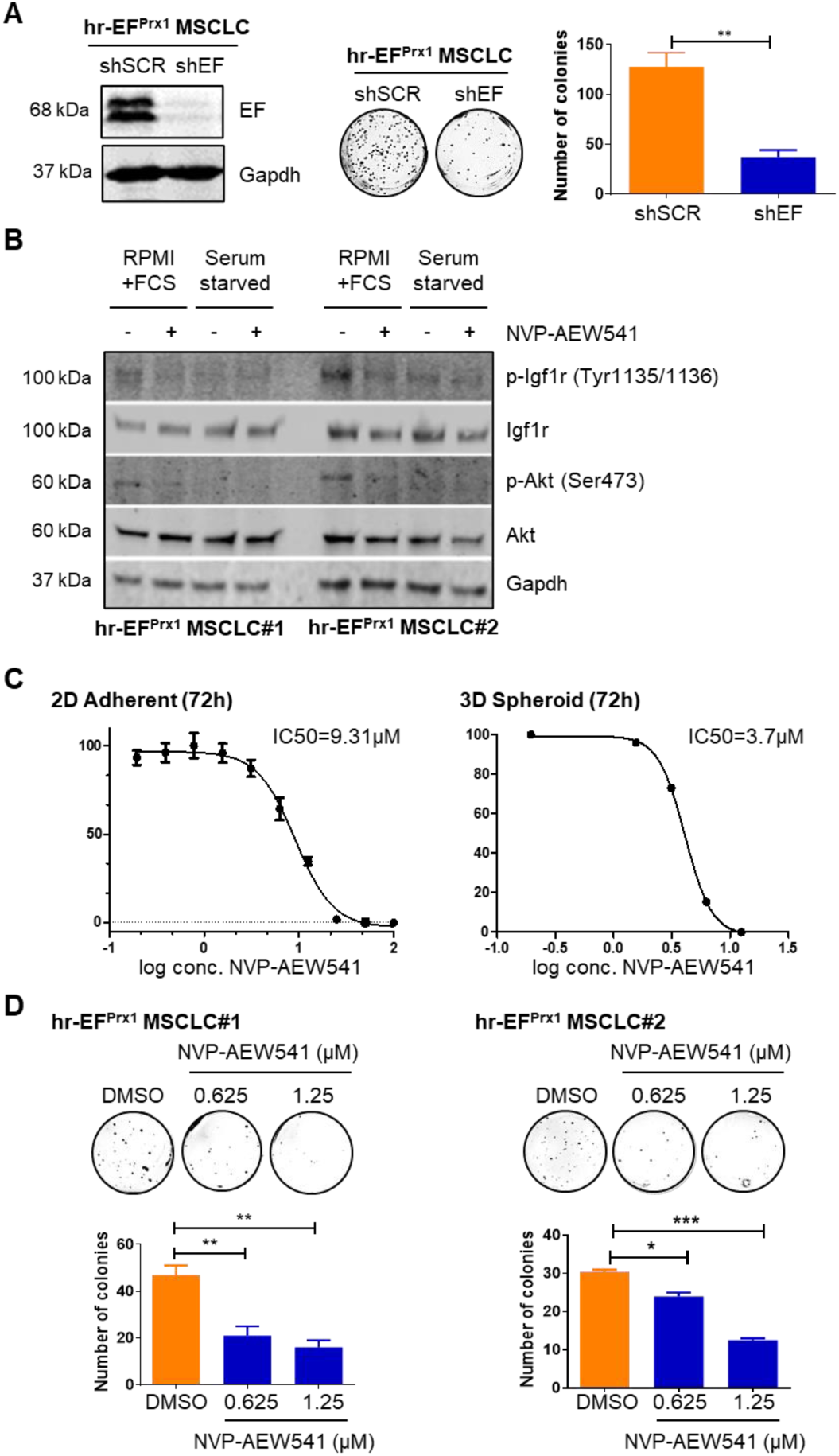
Sustained hr-EF^Prx1^ MSCLC growth depends on EF and paracrine IGF-1 expression. **A** Dependence of hr-EF^Prx1^ MSCLC on continuous EF expression. Left: Western blot analysis of EF knockdown efficacy in hr-EF^Prx1^ cells using EF specific shRNA (sh-EF). Scrambled shRNA was used as a non-targeting control (sh-scr). Middle: representative images of soft agar colony assay for cells transduced with the indicated shRNAs. Right: Quantification of soft agar colonies. Statistics were calculated by two-sided, unpaired Student’s t-test. **B** IGF-1R and Akt phosphorylation status of hr-EF^Prx1^ MSCLC propagated in growth medium supplemented with 10%FCS or serum-starved for 5 hours, followed by treatment with the IGF-1R inhibitor NVP-AEW541 for 2 hours at 1µM (compared to non-treated control). **C** Determination of IC50 values for NVP-AEW541 on hr-EF^Prx1^ MSCLCs in both 2D and 3D (spheroid) formats. **D** Representative pictures and quantification of soft agar colony formation of hr-EF^Prx1^ cells after 12 days incubation in absence (DMSO) or presence of NVP-AEW541 with inhibitor replenishment every 3 days. Colonies > 0.5 mm were counted using ImageJ software. Data are expressed as mean +/- SD. from three independent experiments, two-way Anova was used to determine statistical significance (*P<0.05, **P<0.01).

**Supplementary Figure S5.**
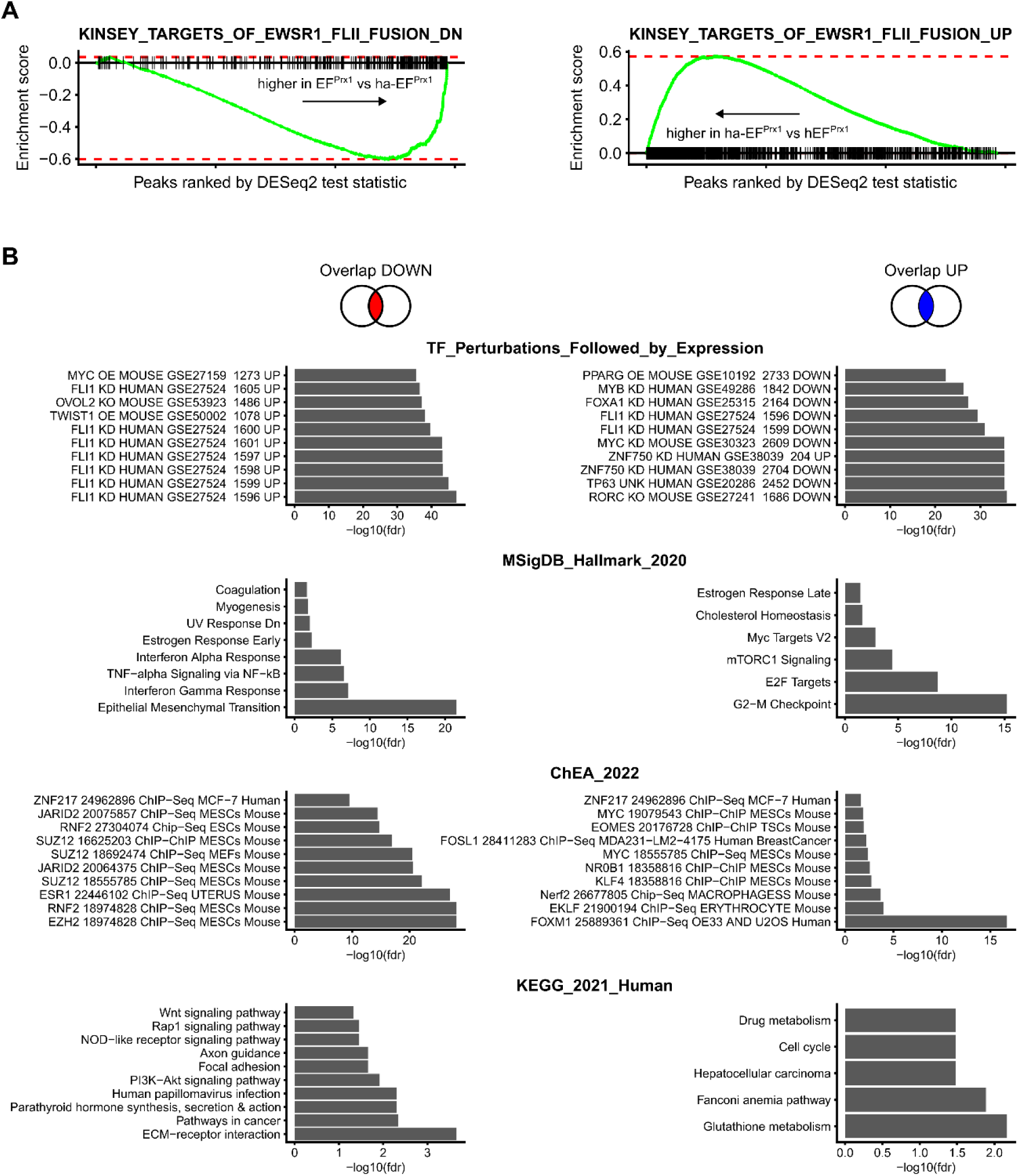
IGF-1 activation and reprogramming enforces an EWS::FLI1 transcriptional signature. **A** Fast gene set enrichment analysis (fGSEA ^32^) of genes down-and up-regulated by the EWS::FLI1 fusion ^91^ and genes up-/down-regulated in ha-EF^Prx1^ MSCLCs. Genes were pre-ranked by the DESeq2 test statistic for the comparison of ha-EF^Prx1^ MSCLC vs. EF^Prx1^ MSCLCs. **B** Plots showing functional gene annotations overrepresented among down-regulated (left) and up-regulated (right) genes in ha-/hr--EF^Prx1^ MSCLCs (overlaps from Fig. 4B). Each plot panel lists the top 10 terms (ranked by number of overlapping genes) and each bar shows the negative logarithm of the FDR-corrected P-value. Only significant enrichments are shown (P_adj_ <= 0.05). Enrichment was calculated using hypeR ^89^. See Table S8.

**Supplementary Figure 6.**
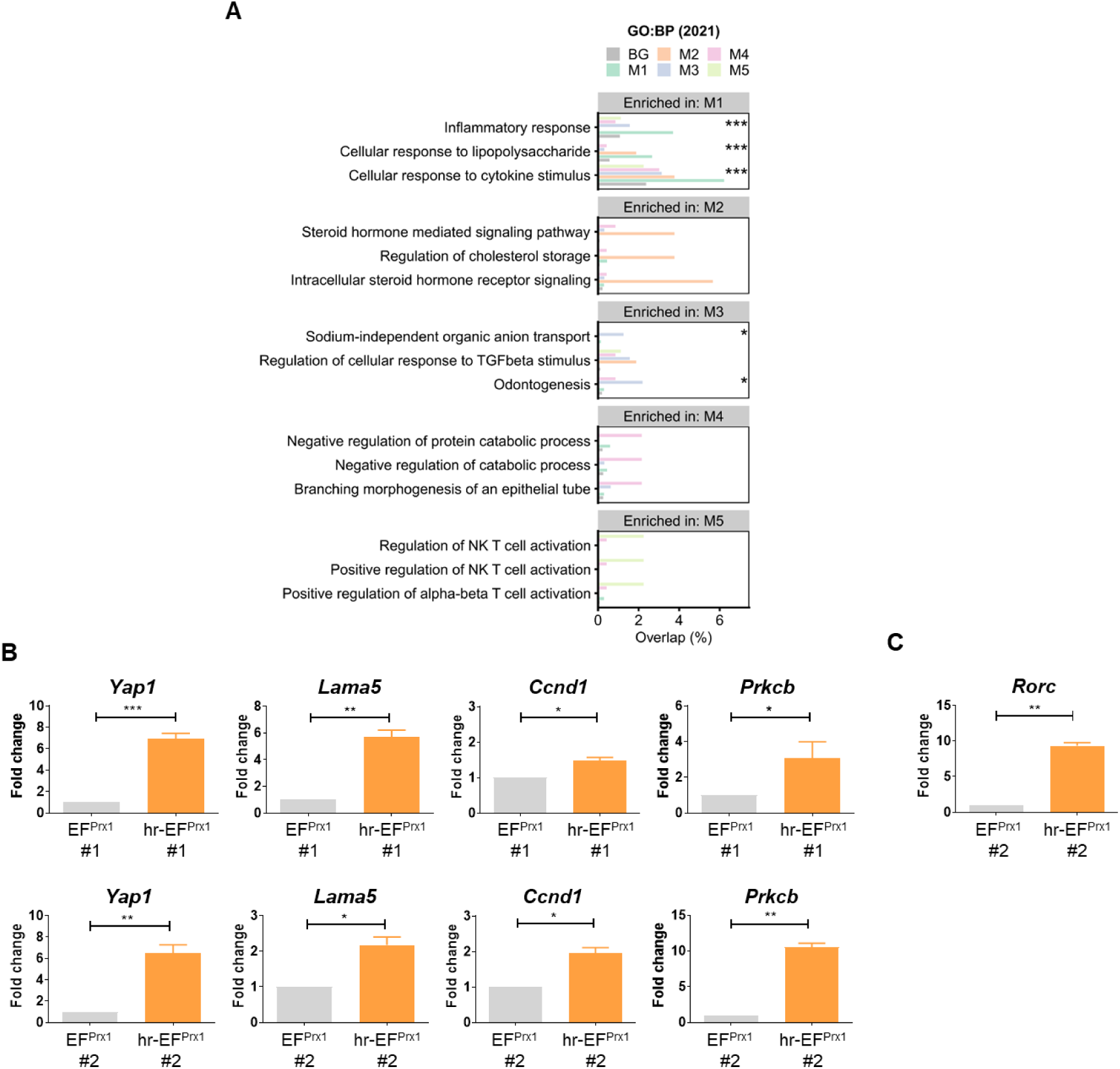
Characterization of chromatin cluster module M2. **A** Plots showing gene ontology (biological process) terms overrepresented for genes associated with peaks (nearest gene) belonging to each of the five chromatin accessibility clusters displayed in Figure 5A. Each plot panel lists the top 3 terms per module, and each bar indicates the percentage of peaks with at least one match to the given term. Enrichment was calculated using hypeR ^89^. *, P_adj_ < 0.05; **, P_adj_ <= 0.01; ***, P_adj_ <= 0.005. See Table S9. **B** Gene expression changes of genes linked to chromatin cluster module M2. Quantitative analysis by RT-qPCR of relative mRNA expression levels of (from left to right) *Yap1, Lama5, Prkcb, and Ccnd1* in hr-EF^Prx1^ MSCLC *versus* EF^Prx1^ MSCLC. Statistics were calculated by one-sided, paired Student’s t-test. Data is presented as mean ± SE (n = 3), ***P<0.001 **P<0.005. *P<0.01. **C** Quantitative analysis by measuring the relative mRNA expression levels of *RorC* for parental and hr-EF^Prx1^ MSCLC#1 and #2. Statistics were calculated by one-sided, paired Student’s t-test. Data is presented as mean ± SE (n = 3), **P<0.005.

**Supplementary Figure 7.**
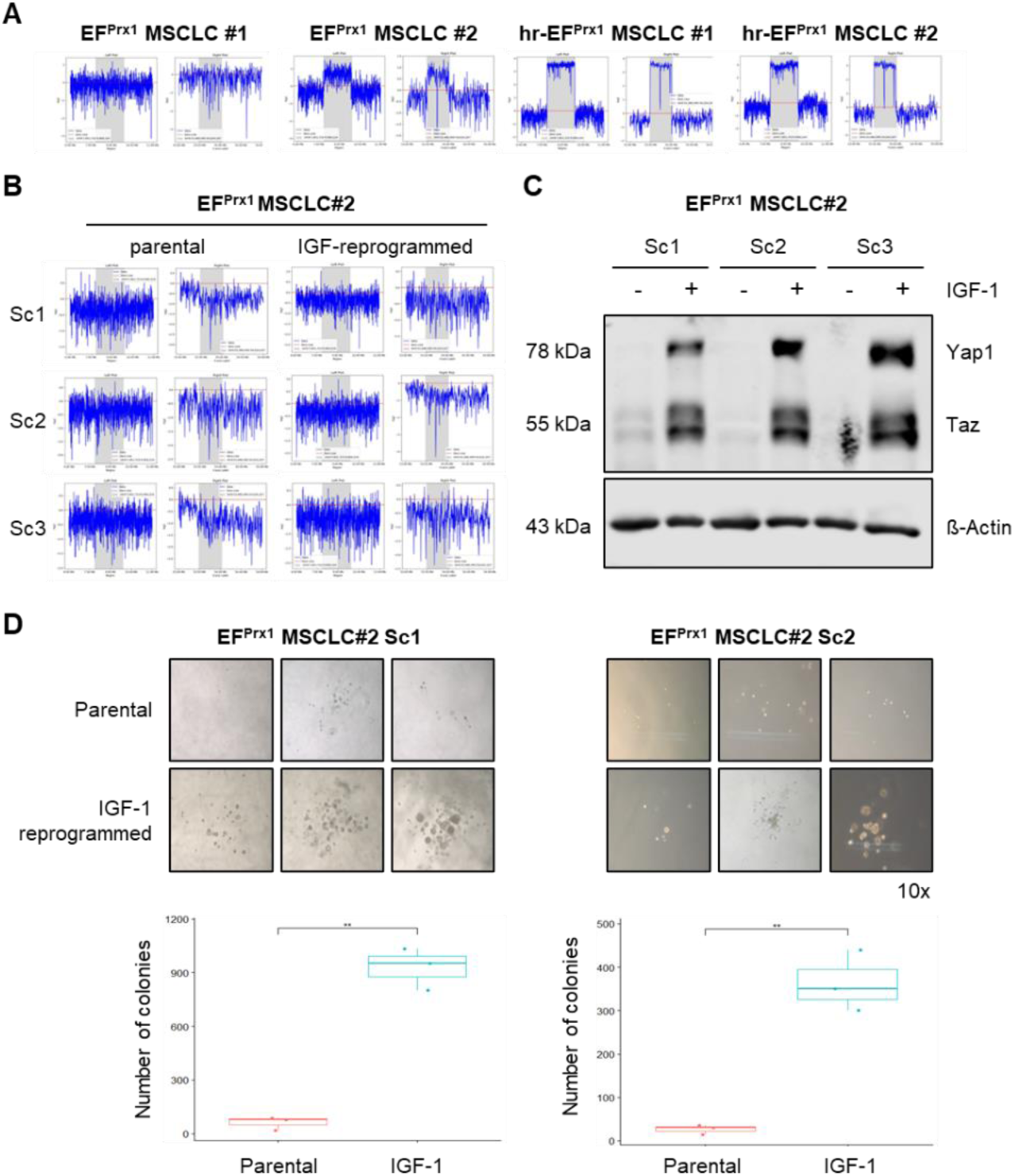
Yap1 induction in IGF-1 reprogrammed cells is independent of but supported by sub-clonal copy number gains at chromosome 9q. **(A)** LcWGS profiles of chromosome 9q regions chr9:7,851,723-9,900,224 (left) and chr9:53,480,495-54,024,207 (right) (grey shaded areas) for bulk parental and IGF-1 reprogrammed derivative EF^Prx1^ MSCLC pools #1 and #2. **(B)** Corresponding lcWGS profiles for hr-EF^Prx1^ MSCLC #2 derived single clones sc1-3. Log2-fold copy number change (y-axis) is presented at two scales for each condition (left and right plots). **(C)** Immuno-blot analysis of Yap1, Taz, and beta actin protein expression in untreated and IGF-1 reprogrammed single-cell clones sc1-3 derived from parental EF^Prx1^ MSCLC#2. **(D)** Soft-agar colony formation of parental and hr-EF^Prx1^ single cell clones sc1 and sc2.

**Supplementary Figure 8.**
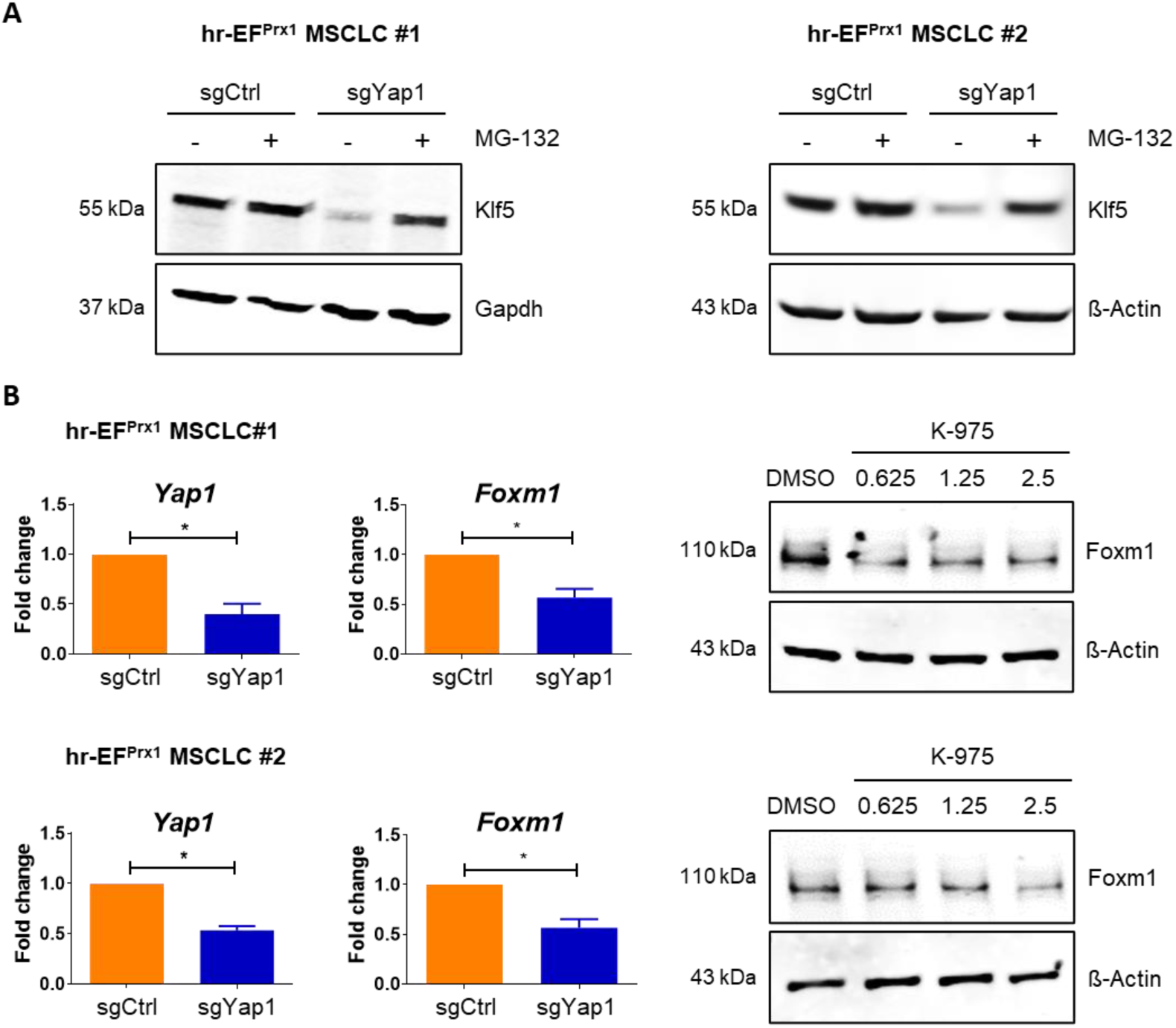
Yap1 knockdown results in diminished FoxM1 expression. **A** Klf5 protein expression in hr-EF^Prx1^ MSCLC upon knockout of Yap1 in presence or absence of the proteasome inhibitor MG135 ( 5 µM, for 5 hr). **B** Left panel: Quantitative analysis of Yap1 expression by RT-qPCR upon sgRNA mediated knockout of Yap1. Statistics were calculated by one-sided, paired Student’s t-test. Data is presented as mean ± SE (n = 3), * <0.05. Right panel: Immunoblot analysis of FoxM1 expression in hr-EF^Prx1^ MSCLC#2 upon treatment with increasing concentrations of the Tead palmitoylation pocket inhibitor K-975 for 24 hr.

## Supplementary Tables

**Supplementary Table S1.** Overview of RNA-seq and ATAC-seq datasets used in this study

**Supplementary Table S2.** Differential gene expression analysis EFPrx1 vs Wt MSCLC

**Supplementary Table S3.** Ontology enrichments for up-and down-regulated genes in EFPrx1 MSCLC

**Supplementary Table S4.** ATAC-seq peaks

**Supplementary Table S5.** Differential accessibility analyses (ATAC-seq)

**Supplementary Table S6.** Differential gene expression analysis ha-EFPrx1 vs EFPrx1 MSCLC

**Supplementary Table S7.** Differential gene expression analysis hr-EFPrx1 vs EFPrx1 MSCLC

**Supplementary Table S8.** Ontology enrichments for up-and down-regulated genes in ha/hr-EFPrx1 MSCLC

**Supplementary Table S9.** Gene ontology and motif enrichment results for ATAC-seq peak modules

