## Supplementary Figures for "YAP1 is a key regulator of EWS::FLI1-dependent malignant transformation upon IGF-1 mediated reprogramming of bone mesenchymal stem cells"

### Slide 1
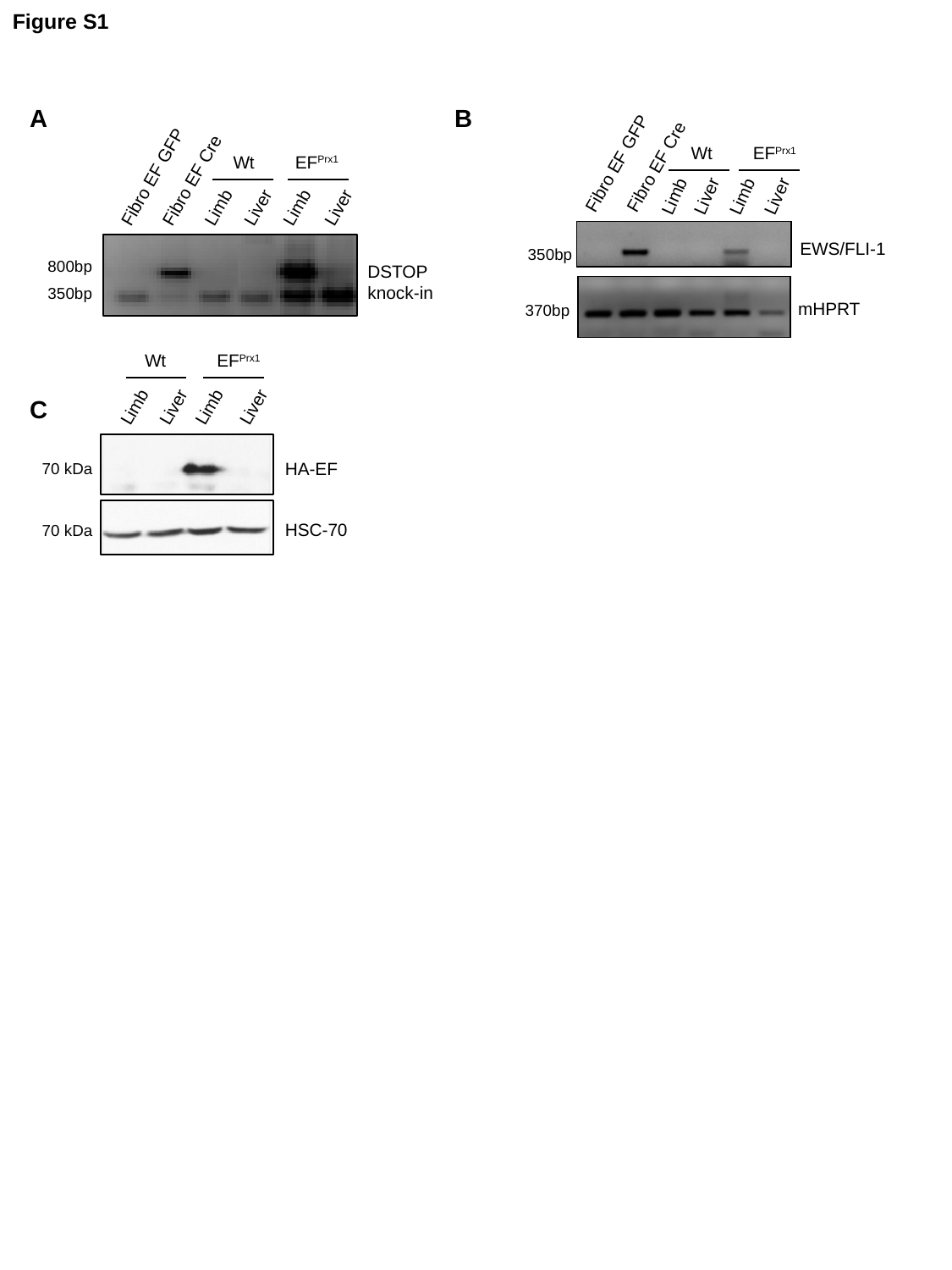

Figure S1
Wt
EF^Prx1^
Fibro EF GFP
Fibro EF Cre
Limb
Limb
Liver
Liver
EWS/FLI-1
350bp
mHPRT
370bp
A
B
Wt
EF^Prx1^
Fibro EF GFP
Fibro EF Cre
Limb
Liver
Limb
Liver
800bp
DSTOP
knock-in
350bp
Wt
EF^Prx1^
C
Limb
Liver
Limb
Liver
HA-EF
70 kDa
HSC-70
70 kDa

### Slide 2
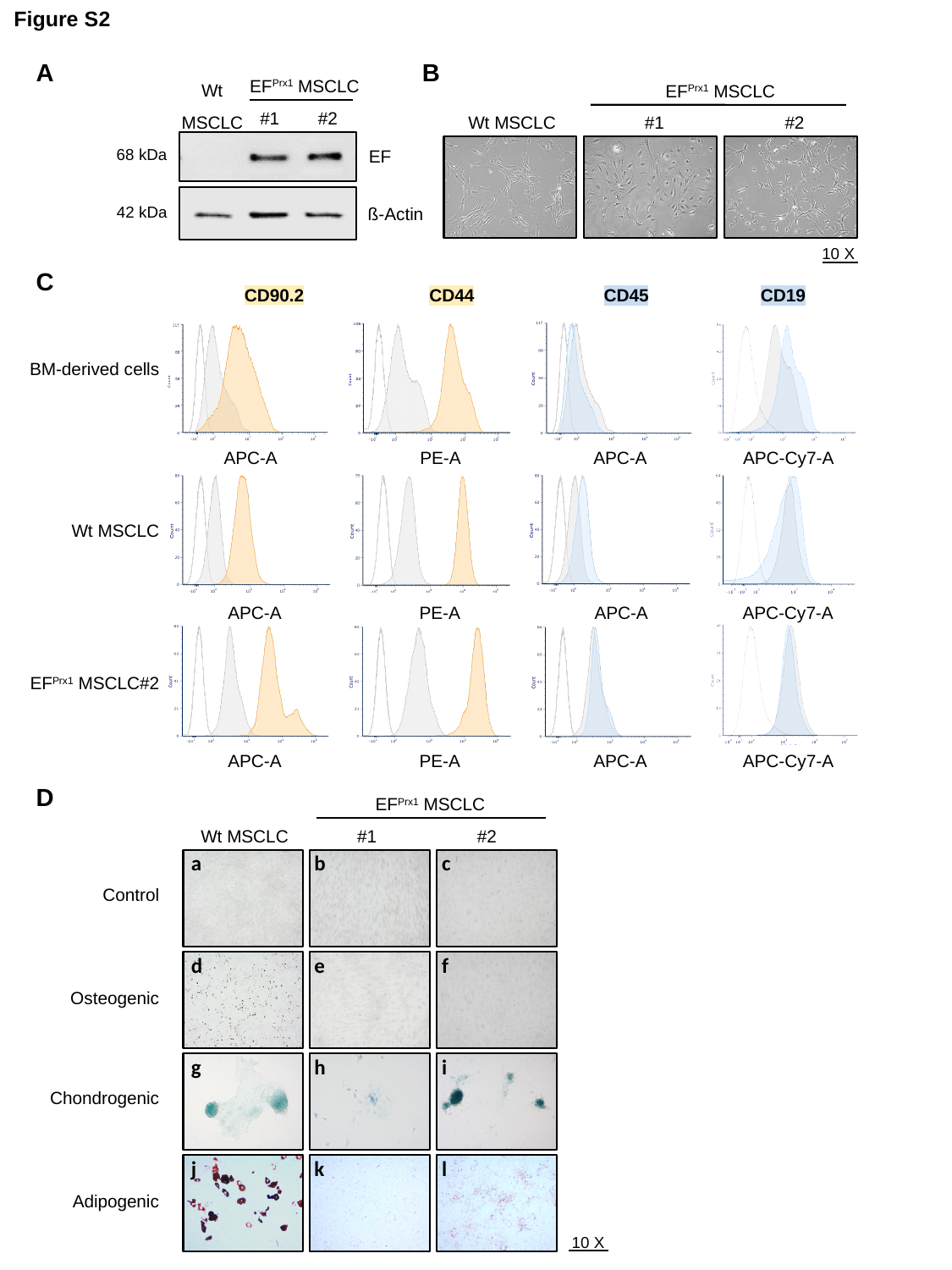

Figure S2
A
B
Wt MSCLC
EF^Prx1^ MSCLC
EF^Prx1^ MSCLC
#1
68 kDa
EF
42 kDa
ß-Actin
#2
Wt MSCLC
#2
#1
10 X
C
CD90.2
CD44
CD45
CD19
BM-derived cells
APC-A
PE-A
APC-A
APC-Cy7-A
Wt MSCLC
APC-A
PE-A
APC-A
APC-Cy7-A
EF^Prx1^ MSCLC#2
APC-A
PE-A
APC-A
APC-Cy7-A
D
EF^Prx1^ MSCLC
Wt MSCLC
#1
#2
a
b
c
Control
d
e
f
Osteogenic
g
h
i
Chondrogenic
j
k
l
Adipogenic
10 X

### Slide 3
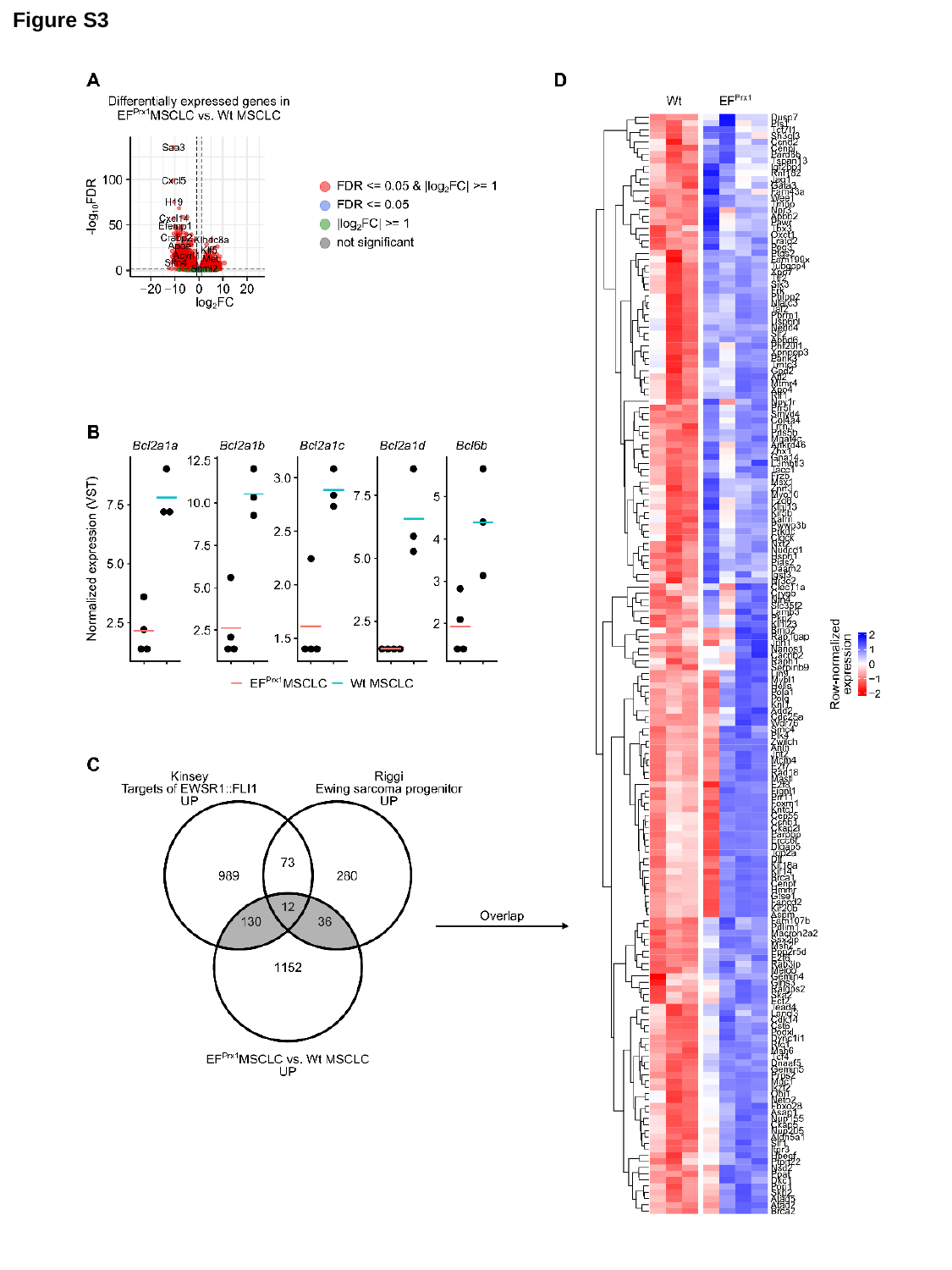

Figure S3

### Slide 4
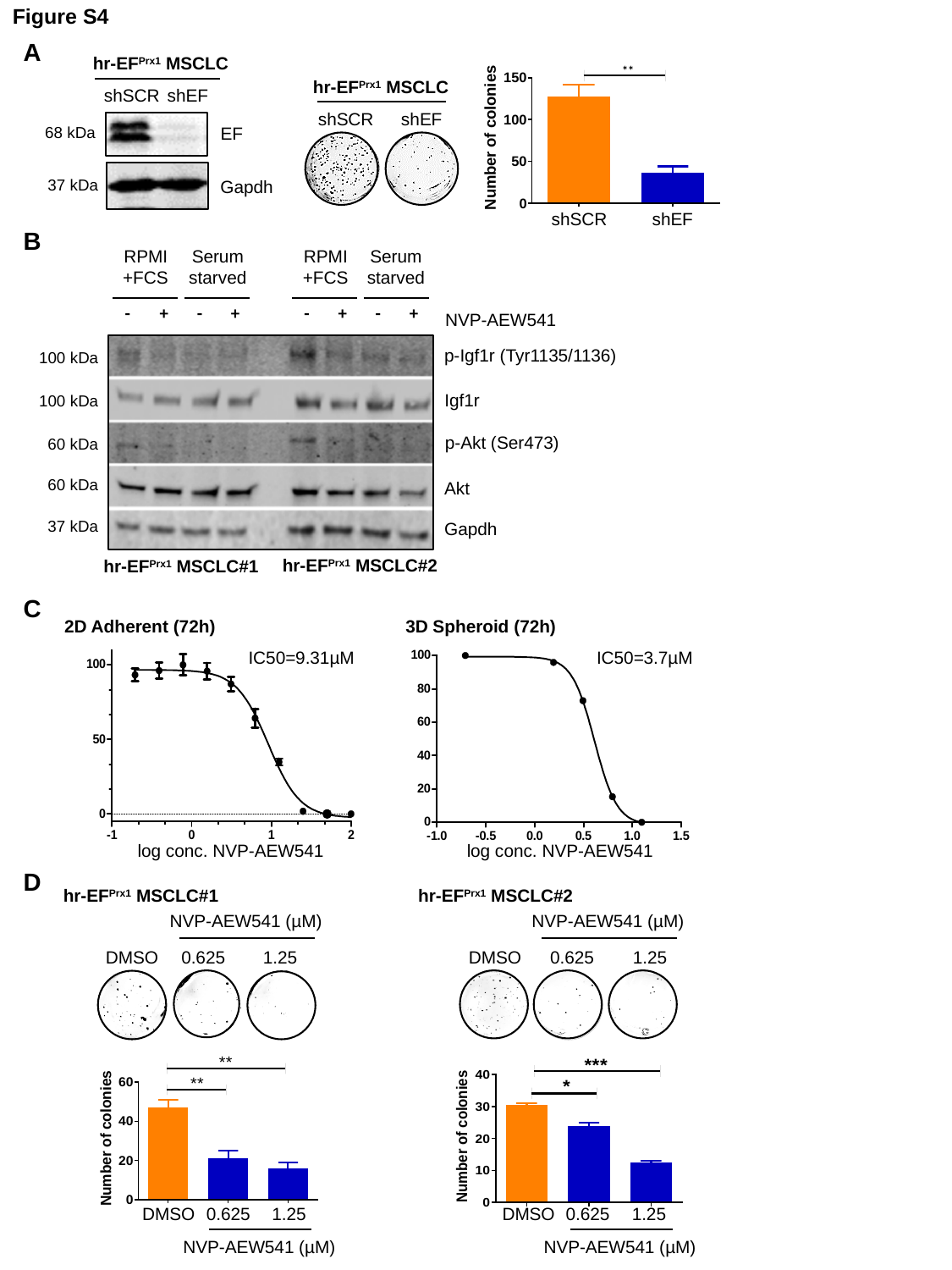

Figure S4
A
hr-EF^Prx1^ MSCLC
shSCR
shEF
EF
68 kDa
37 kDa
Gapdh
hr-EF^Prx1^ MSCLC
shSCR
shEF
shSCR
shEF
B
RPMI+FCS
Serum starved
RPMI+FCS
Serum starved
| - | + | - | + | | - | + | - | + |
| --- | --- | --- | --- | --- | --- | --- | --- | --- |
NVP-AEW541
p-Igf1r (Tyr1135/1136)
100 kDa
Igf1r
100 kDa
p-Akt (Ser473)
60 kDa
60 kDa
Akt
37 kDa
Gapdh
hr-EF^Prx1^ MSCLC#2
hr-EF^Prx1^ MSCLC#1
C
2D Adherent (72h)
3D Spheroid (72h)
IC50=9.31µM
IC50=3.7µM
log conc. NVP-AEW541
log conc. NVP-AEW541
D
hr-EF^Prx1^ MSCLC#1
hr-EF^Prx1^ MSCLC#2
NVP-AEW541 (µM)
NVP-AEW541 (µM)
DMSO
0.625
1.25
DMSO
0.625
1.25
DMSO
0.625
1.25
DMSO
0.625
1.25
NVP-AEW541 (µM)
NVP-AEW541 (µM)

### Slide 5
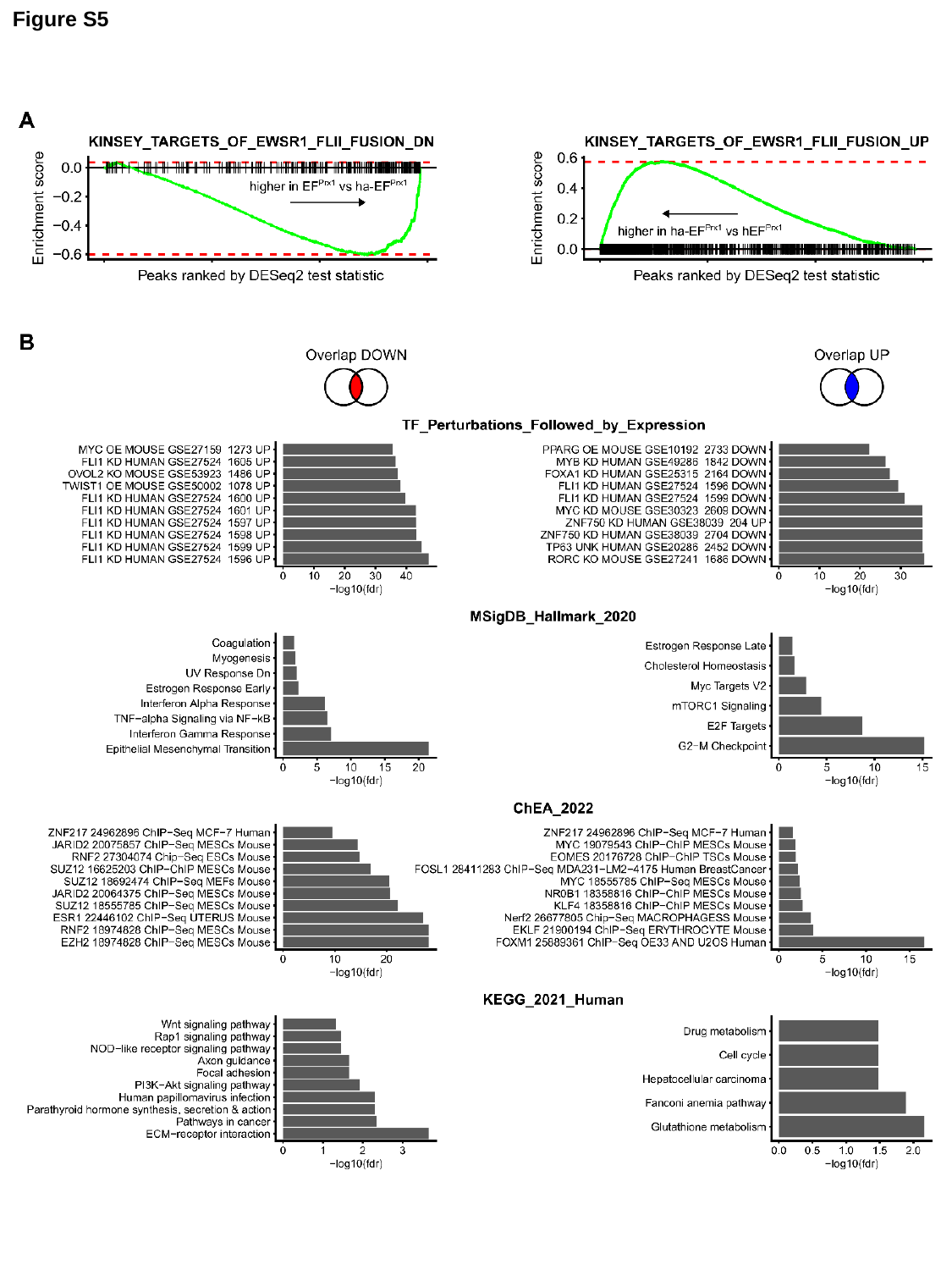

Figure S5

### Slide 6
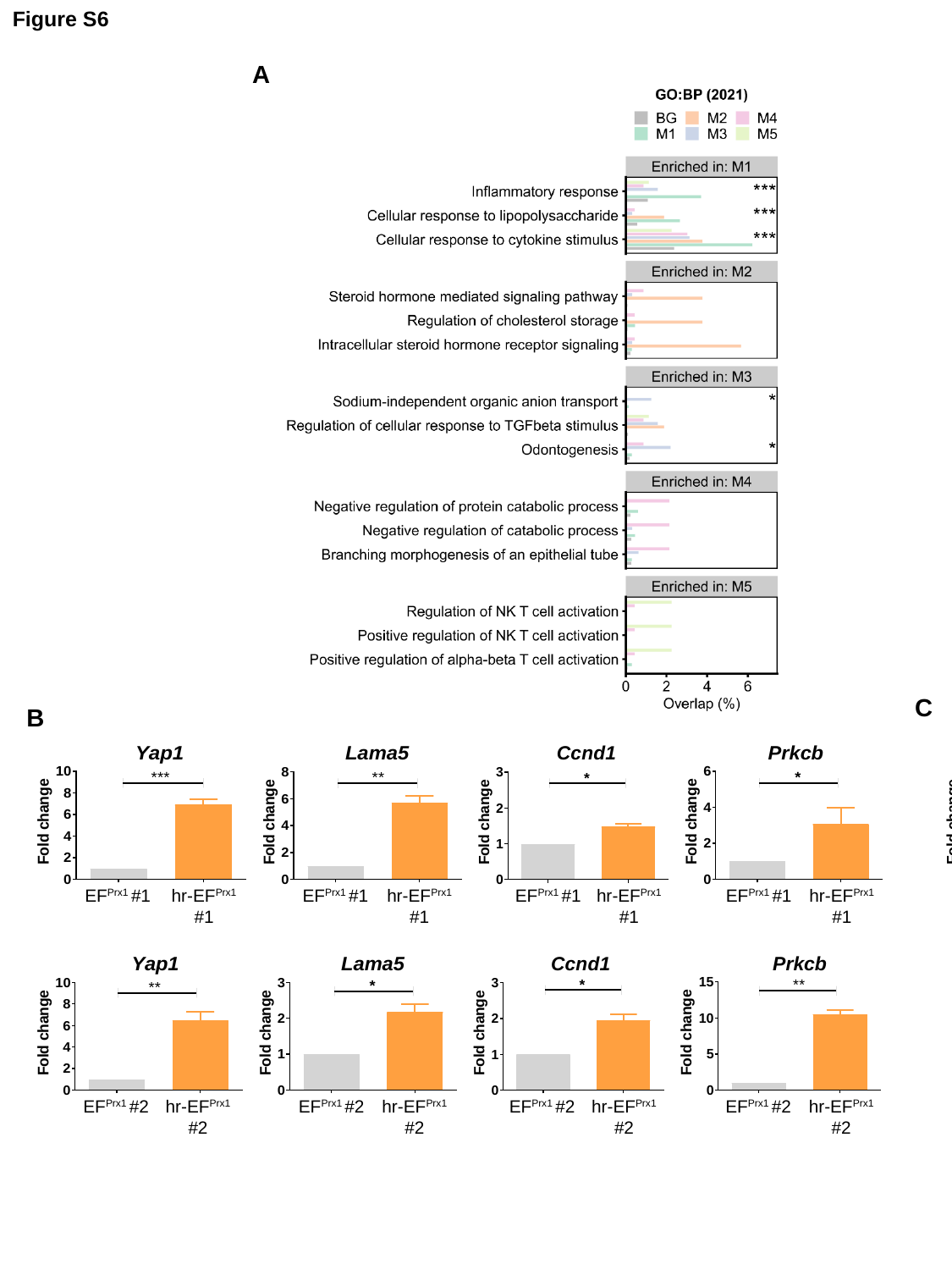

Figure S6
A
C
EF^Prx1^ #2
hr-EF^Prx1^ #2
B
EF^Prx1^ #1
hr-EF^Prx1^ #1
EF^Prx1^ #1
hr-EF^Prx1^ #1
EF^Prx1^ #1
hr-EF^Prx1^ #1
EF^Prx1^ #1
hr-EF^Prx1^ #1
EF^Prx1^ #2
hr-EF^Prx1^ #2
EF^Prx1^ #2
hr-EF^Prx1^ #2
EF^Prx1^ #2
hr-EF^Prx1^ #2
EF^Prx1^ #2
hr-EF^Prx1^ #2

### Slide 7
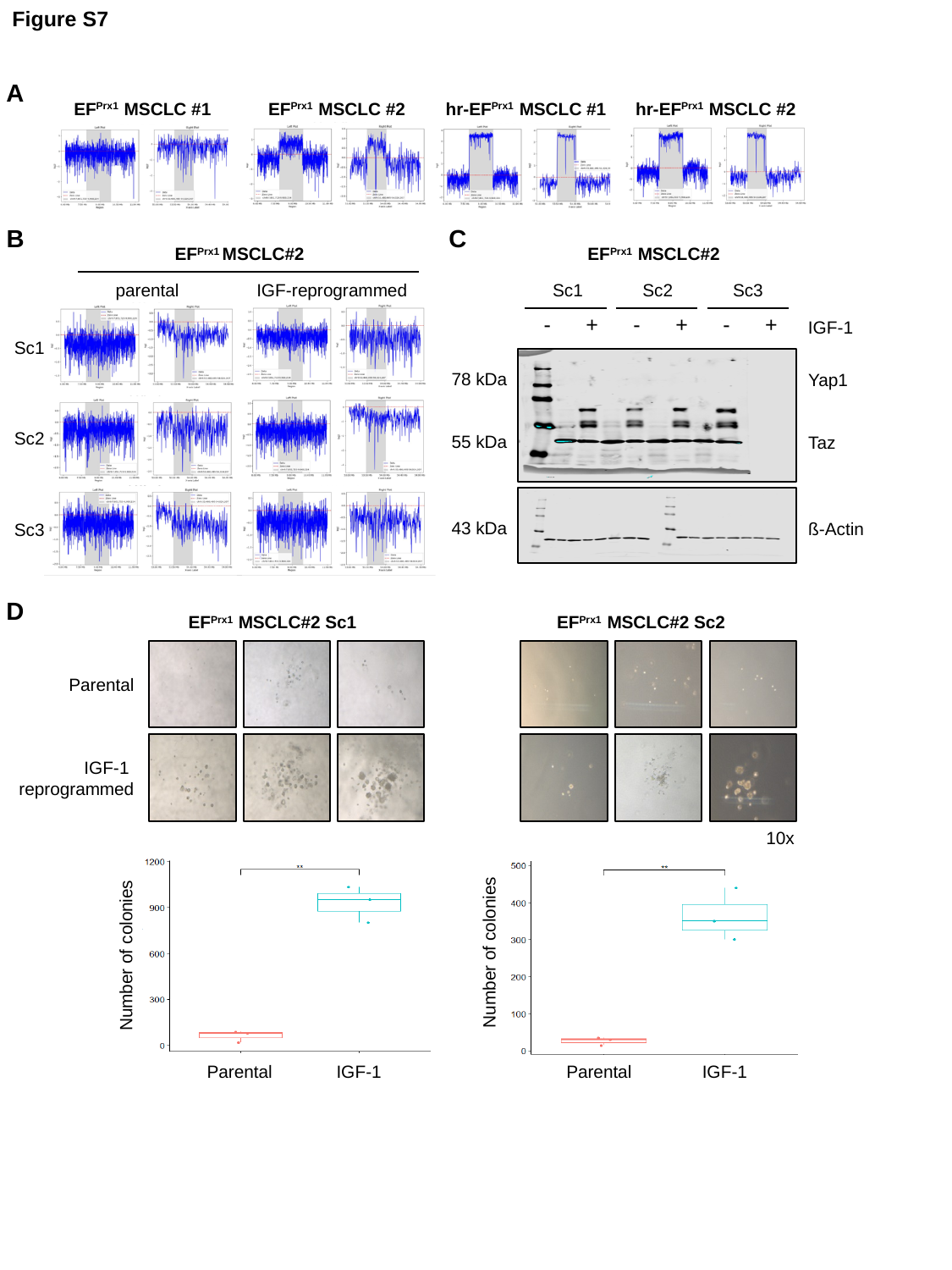

Figure S7
A
EF^Prx1^ MSCLC #1
EF^Prx1^ MSCLC #2
hr-EF^Prx1^ MSCLC #1
hr-EF^Prx1^ MSCLC #2
B
C
EF^Prx1^ MSCLC#2
EF^Prx1^ MSCLC#2
parental
IGF-reprogrammed
Sc1
Sc2
Sc3
| - | + | - | + | - | + |
| --- | --- | --- | --- | --- | --- |
IGF-1
Sc1
78 kDa
Yap1
Sc2
55 kDa
Taz
43 kDa
ß-Actin
Sc3
D
EF^Prx1^ MSCLC#2 Sc1
EF^Prx1^ MSCLC#2 Sc2
Parental
IGF-1
reprogrammed
10x
Number of colonies
Number of colonies
Parental
IGF-1
Parental
IGF-1

### Slide 8
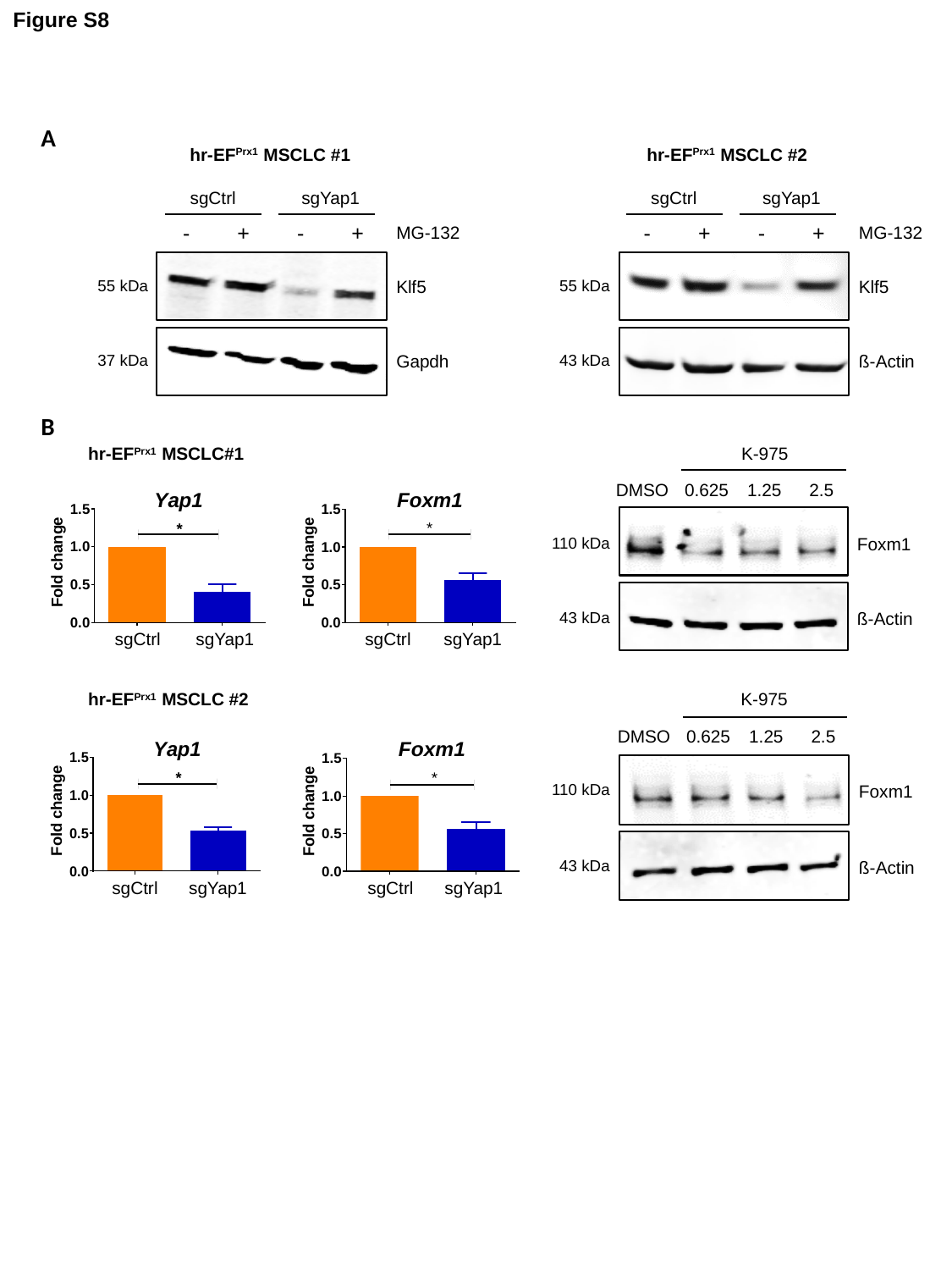

Figure S8
A
hr-EF^Prx1^ MSCLC #1
hr-EF^Prx1^ MSCLC #2
sgCtrl
sgYap1
sgCtrl
sgYap1
| - | + | - | + |
| --- | --- | --- | --- |
MG-132
| - | + | - | + |
| --- | --- | --- | --- |
MG-132
55 kDa
Klf5
55 kDa
Klf5
37 kDa
Gapdh
43 kDa
ß-Actin
B
hr-EF^Prx1^ MSCLC#1
K-975
DMSO
0.625
1.25
2.5
110 kDa
Foxm1
43 kDa
ß-Actin
sgCtrl
sgYap1
sgCtrl
sgYap1
hr-EF^Prx1^ MSCLC #2
K-975
DMSO
0.625
1.25
2.5
110 kDa
Foxm1
43 kDa
ß-Actin
sgCtrl
sgYap1
sgCtrl
sgYap1
